# Multi-lingual multi-platform investigations of online trade in jaguar parts

**DOI:** 10.1101/2022.09.19.508455

**Authors:** John Polisar, Charlotte Davies, Thais Morcatty, Mariana da Silva, Song Zhang, Kurt Duchez, Julio Madrid, Ana Elisa Lambert, Ana Gallegos, Marcela Delgado, Ha Nguyen, Robert Wallace, Melissa Arias, Vincent Nijman, Jon Ramnarace, Roberta Pennell, Yamira Novelo, Damian Rumiz, Kathia Rivero, Yovana Murillo, Monica Nunez Salas, Heidi Kretser, Adrian Reuter

## Abstract

We conducted research to understand online trade in jaguar parts and develop tools of utility for jaguars and other species. Our research took place to identify potential trade across 31 online platforms in Spanish, Portuguese, English, Dutch, French, Chinese, and Vietnamese. We identified 230 posts from between 2009 and 2019. We screened the images of animal parts shown in search results to verify if from jaguar; 71 posts on 12 different platforms in four languages were accompanied by images identified as definitely jaguar, including a total of 125 jaguar parts (50.7% posts in Spanish, 25.4% Portuguese, 22.5% Chinese and 1.4% French). Search effort varied among languages due to staff availability. Standardizing for effort across languages by dividing number of posts advertising jaguars by search time and number individual searches completed via term/platform combinations, the adjusted rankings of posts were: Portuguese #1, Chinese 2 (time) & 3 (searches), Spanish 3 & 4; French 5 & 4; English 5 & 2, and Dutch 6. Teeth were most common; 156 posts offered at least 367 apparent teeth. From these, 95 teeth were assessed as definitely jaguar; 71 jaguar teeth could be linked to a location, with the majority of the 71 offered for sale from Mexico, China, Bolivia, and Brazil (26.8, 25.4, 16.9, and 12.7% respectively). Ranking of number of teeth was Mexico (19), China (18), Bolivia (12), Brazil (9), Peru/Ecuador (most accurate probable location) (8), Venezuela (3), Guadeloupe (1), and Uruguay (1). The second most traded item, skins and derivative items were only identified from Latin America: Brazil (7), was followed by Peru (6), Bolivia (3), Mexico (2 and 1 skin piece), and Nicaragua and Venezuela (1 each). Whether by number of posts or pieces, the ranking of parts was teeth, skins/pieces of skins, heads, and bodies. Our research presents a snapshot of online jaguar trade and methods that may have utility for many species now traded online. Our research took place within a longer-term project to assist law enforcement in host countries to better identify potential illegal trade online, with research findings informing hubs in Latin America for building such capacity.

## Introduction

In the first High-Level Conference on Illegal Wildlife Trade in the Americas in 2019, the jaguar (*Panthera onca*) was declared an emblematic species of the Americas [1] and as a symbol of the struggle against illegal wildlife trade. This physically impressive keystone and umbrella species had a historic range that extended from the southwestern United States to the middle of Argentina [2,3,4,5]. The historic range has been diminished by over 50% in the last 100 years, due to factors that include habitat loss, prey depletion, and direct killing and trade in jaguar parts [3]. Recent population estimates vary between 64,000 [6] and 173,000 [7]. Rapid rates of habitat loss [8], combined with the small scale and frequent biases of jaguar population sampling sites when compared to the species’ vast and variable range, and the corresponding high degrees of uncertainty, suggest that erring towards the more conservative estimate may be advisable.

Although the current IUCN Red List Status of the jaguar is Near Threatened (NT) [3], finer-grained analyses have revealed that the majority of sub-populations are endangered (E), and some are critically endangered (CR) [6,9]. The eighteen countries where jaguar populations currently occur include: Argentina; Belize; Bolivia; Brazil; Colombia; Costa Rica; Ecuador; French Guiana; Guatemala; Guyana; Honduras; Mexico; Nicaragua; Panama; Paraguay; Peru; Suriname; and Venezuela [4]. The jaguar is threatened by trade in body parts, with potentially severe impacts, similar to trade in Asia of tigers and other big cats [9]. There are multiple demand-drivers for big cat parts in Asia, including but not limited to teeth and claws for jewelry and status symbols, bones for medicines and prestige tonics, meat for luxury dishes, and skins for luxury décor, taxidermy and other uses [10,11,12]. The drivers behind trade in jaguar parts are at least as, if not more, diverse and complex [9,13]. Trade in big cat parts has had such drastic impacts on native species in Asia that preliminary evidence of East-West links in trade in jaguar parts, generated concern that a similar surge of trade in jaguar parts could generate catastrophic declines in the New World, and stimulated investigations at multiple scales [9,14,15,16,17,18,19,20,21]. Their findings to-date have included recognition of notable deficiencies in the data needed to understand and disrupt the illegal trade, exacerbated by inconsistent and often weak law enforcement [9,19].

In a broader context, Illegal Wildlife Trade (IWT) - defined as the supplying, selling, purchasing or transport of wildlife and wildlife parts and products in contravention of national or international laws or treaties - has become the primary threat to survival for many species in the New World (Reuter et al. 2018), and has driven many species to near extinction. Such is the case with the nearly extinct vaquita (*Phocoena sinus*), an indirect victim of illegal take of the totoaba fish (*Totoaba macdonaldi*), whose swim bladder is prized in Asia; Spix’s macaw (*Cyanpositta spixii*) from BraziI, considered as possibly extinct in the wild; and a number of freshwater turtle species whose populations have plummeted due to illegal international trade [22,23,24,25,26,27,28,29]. Awareness, understanding, and national enforcement capacity and actions - all need to increase to counter these trends.

Trade in jaguar parts is not new; but currently combined with two centuries of rapidly increasing levels of habitat loss, and human-jaguar conflict, it has the potential to be a serious threat to the species’ persistence. Archaeological records indicate jaguar body parts traveled long distances across the Caribbean Sea as early as the Ceramic Age (500 BC to AD 1500) potentially transported as prized items of exchange between Amerindian and Caribbean societies [30]. In 1876, a Cheyenne warrior wore a mysterious spotted robe and belt during the battle of the Little Bighorn in Montana more than 1200 km north of jaguar range [31]. As human populations grew and resource extraction escalated during the European colonization of the Americas, commerce in jaguar parts increased. During the 18th century, there are records of approximately 2,000 jaguars being exported annually from Buenos Aires to Europe for the fur industry [32]. Jaguar trade reached unprecedented commercial levels during the first three-quarters of the 20th century, when the fashion for spotted cat furs reached its peak. In Brazil, an estimated 180,000 jaguars were killed during this period [33], causing a widespread population decline. In response to the imminent extinction risk posed to jaguars and other spotted cats by the fur trade, in 1975, the Convention on International Trade in Endangered Species of Wild Fauna and Flora (CITES) listed these species under Appendix I, prohibiting their commercial trade across international boundaries [15,34]. By the end of the 20th century, international jaguar trade was virtually over [9,33,35,36]. Jaguar killing and trading continued domestically at much lower intensity, mostly as an opportunistic response to jaguar-livestock conflicts, preventative killing, and fear [17, 37].

Since 2010, but increasingly from 2014 onwards, an apparent resumption of trade in jaguar parts raised concerns about the characteristics and scale of the commerce and its potential effects on jaguar populations. These reports focused on Bolivia and Suriname [4,14,16,38,39,40]. Between 2014 and 2020, 22 of 52 jaguar trafficking events in Bolivia were directly related to China (42%) either as people of Chinese origin living in Bolivia involved as some part of the supply chain (sellers, collectors/middle persons, buyers), or geographically, through seized packages destined for China, or seizures in China of Bolivian origin. Although only 42% of all detected jaguar trafficking events, these 22 events accounted for 642 of 673 (95.39%) seized jaguar canines. Similar cases were registered in Suriname, including the sale of jaguar body parts in physical and online markets, and the preparation of jaguar paste [16,40,41]. The extent to which trade in jaguar parts is related to increased Chinese presence and investments in Latin America has been explored [19]. In some isolated cases, those links have seemed clear, but the situation has required more holistic investigations.

The most worrisome element of the apparent resurgence has been its potential to derail a post-CITES listing jaguar recovery underway in some areas, and exacerbate declines in others. Massive losses of jaguar habitat range wide imply rapid declines in the global jaguar population [42], yet stability has been achieved in some strongholds. Although these examples of conservation success tend to be limited in spatial extent, jaguar population stability, and in some cases increases, are evident in some areas and are linked to proven conservation interventions [43,44,45,46]. Illegal trade threatens those advances across jaguar range. Given how illegal trade drove tigers in Asia to near extinction [47], it is imperative that we act promptly to better understand and address this similar threat for jaguars. Methods to monitor IWT and identify junctures where it can be disrupted have primarily focused on: i) direct observation/physical examinations of material being shipped, or sold in markets; ii) official data resulting from enforcement actions, such as seizures, and; iii) advertisements using traditional channels through open sources or specialized group publications, discussion groups and hobbyist and collector events [9,48,49]. Yet like other crimes, IWT is growing in its level of sophistication, and due to technological advances and the expansion of the internet, a growing proportion of IWT is now taking place online [50,51,52]. Web platforms have replaced brick and mortar wildlife trade in many countries [53]. From a law enforcement perspective, specialized investigative techniques typically used to address other types of illicit trade have rarely been applied, including controlled deliveries, electronic surveillance, undercover infiltration of criminal enterprises, capture of digital evidence, deployment of tracking devices within contraband shipments, or online searches [54]. In particular, the application of online monitoring of wildlife trade is a young field [49,52,55,56], with few methodological recommendations and guidance available for online monitoring [57]. However, online trade in wildlife has expanded rapidly and tools to investigate and curb it have been urgently needed [58].

To address knowledge gaps about the extent of trade in jaguar parts, particularly about the use of internet in the trade, we designed and conducted research on online trade in jaguar parts and products. Our searches were conducted in every major national language used within jaguar range, as well as Chinese and Vietnamese. By searching online in seven languages, our investigation was less restricted to national boundaries than previous research on trade in jaguar parts [13,14,16,18,20,38], potentially identifying posts from anywhere where the seven languages are employed. The two languages based outside of jaguar range, Chinese and Vietnamese, were informed by previous research indicating the countries to be significant for tiger trade [13,59,60,61,62], yet we recommend further research in additional languages may be undertaken in future.

Specific research questions included: 1) how prevalent is the online trade in jaguar parts, in a number of countries, including within Asia? 2) has trade in jaguar parts been under-detected by previous studies in some countries, and if so which ones? 3) what can be done to better understand and interdict illegal trade in jaguar parts? In our aim of providing a larger-scale perspective about online jaguar trade, we needed to develop and test methods to identify information and sought to address gaps in knowledge related to online trade as an integral part of addressing wider threats to wild jaguars. The research findings we present have implications for future research, policy, and enforcement. The methods that we present have led to concrete combatting wildlife trade actions; have been applied across multiple other illegally traded taxa; and are presented as an additional tool to ensure human security in the context of the presumed wildlife trade origins of the devasting global COVID 19 pandemic.

## Methods

We generated methods for online research. These align with subsequent recommendations made by Stringham et al. 2020 [57].

### Period of research

Our online research to identify openly-available posts indicative of trade in jaguar parts and products took place between May 2019 and March 2020, identifying trade posts on 31 online platforms in seven languages – Spanish, Portuguese, English, Dutch, French, Chinese, and Vietnamese. Timeframe of posts was not designated during searches, but we identified posts from 2009 onwards, representing a band of ten years at the outset of the research.

### Platforms reviewed

We followed the definition of ‘platforms’ as “digital services facilitating interaction between two or more distinct but interdependent sets of users” [63]. There are multiple taxonomies for online platforms [64]. We took a simplified user perspective and classified by the primary user purpose of the platform. Platforms were categorized as general search engines, online marketplaces, video-sharing, social networks (House of Lords Select Committee on European Union, 2016, cited in [65]), and weblog platforms (blogs). For ‘online marketplace’ we included e-commerce sites, sites of auction houses, and classified ad sites. We recognize definitions can be broad and platforms can incorporate multiple or hybrid features and functionalities. Given the online space is constantly developing, better definitions may be available in the future. In addition, since our study, new platforms have been created or expanded numbers of active users including across countries. We did not access the ‘Dark Web’ which previous research identified hosted scant volumes of illegal wildlife [65].

### Bias mitigation

Our research was based on language and geography, so considered potential bias in retrieved search results, and attempted to mitigate platform algorithms based on location and search histories to best accomplish unbiased sampling. Methods included regular cleaning of cookies and iterative development of apt search terms across a suite of languages. We first constructed a very extensive list of words combined into search terms used in each language/country to try to avoid missed sampling and bias, whilst searching within and across national languages. These search terms were the lens through which we viewed online traffic. We do not present the list of over 200 search terms that were used in various languages because that may inform traders on how to evade detection from law enforcement.

The investigations followed two paths: 1) structured searches using the most prevalent search engines in each geography and language, and; 2) more intensive searches within specific platforms. The search terms, search engines, and online platforms used were recorded to assess the most productive combinations. The basic structure of these searches was similar across all geographies and languages to achieve a level of standardization, and the resultant ability to make some cross-geography and cross-language comparisons, however there was variance in research time across the languages due to staff availability.

All searches identified the number of productive returns within the first 100 search results, using structured keyword formulations. The second path additionally involved ‘deep dive’ searching within dominant platforms, informed by the most productive keyword structures. Keywords included all jaguar parts and derivatives along with words or currency symbols indicative of trade. We recorded research effort in terms of a) time spent with each search term/platform combination; and b) the quantity of returns from each search term/platform combination, both for unproductive searches and returns, and for productive returns, with jaguar trade posts being the metric of productive searches. The research was informed by guidance to assess the veracity of posts and images (S1 File).

Despite the search method standardization across the 13 researchers conducting searches, level of effort was inherently uneven among the seven languages included in this analysis. Five people searched in Spanish, with the teams divided by regions, Mexico and Mesoamerica (pooled), Colombia and Ecuador (pooled), Peru, Bolivia, and Venezuela, Argentina, Paraguay. Three people searched in English (Belize, Guyana, other), and one each in Portuguese (Brazil), Chinese, Vietnamese, French (French Guiana, Francophone Caribbean, other), and Dutch (Suriname, other). We designed the online research similarly to how we sample wildlife populations [2,66,67,68] with search term / search / engine platform combinations acting as the lens (observer), and the time spent and the number of searches per combination / language / geography being the metric of effort (similar to km transects walked or trap nights) (S1, S2, S3, and S4 Files). This facilitated standardized comparisons of results by search terms and online platform combinations, across languages and geographies. A strict level of standardization could facilitate constant proportion indices [66] across languages and geographies.

Researchers collaboratively designed a data collection record (S1 File) and a database schema in Microsoft Excel organizing one post per row, containing two ‘species’ columns to note when one post showed more than one type of jaguar or other species part, and quantify such parts. Duplicate posts were identified and screened out to count only one post; duplicates referred to whether the post was text only; image only; video only, and researchers assessed whether the post in its entirety was replicated exactly on the same or another platform.

The total of 579 images of animal parts featured in the searches, including multiple images associated with one post, were reviewed to verify if they were from jaguar. These images (whole skins, skin fragments, teeth, claws, skulls, other bones, and jewelry crafted from teeth and other carnivore parts) were examined by two researchers who were the leading global experts at this task, having worked on high-profile successful prosecutions in Bolivia. The two worked in tandem and visually compared with known samples of jaguar parts from museum specimens, confiscated materials in Bolivia, and published literature with images. We did not employ machine-learning in image classification, but future studies may choose to do so. Each image was classified as 1) definitely jaguar; 2) maybe jaguar, ambiguous; 3) definitely not jaguar, with the verification results included in the analysis, against the post per row. The data was reviewed, categorized and cleaned in a progressive triage process and analysis undertaken to produce these results (Fig 1, S2 File).

**Fig 1.**
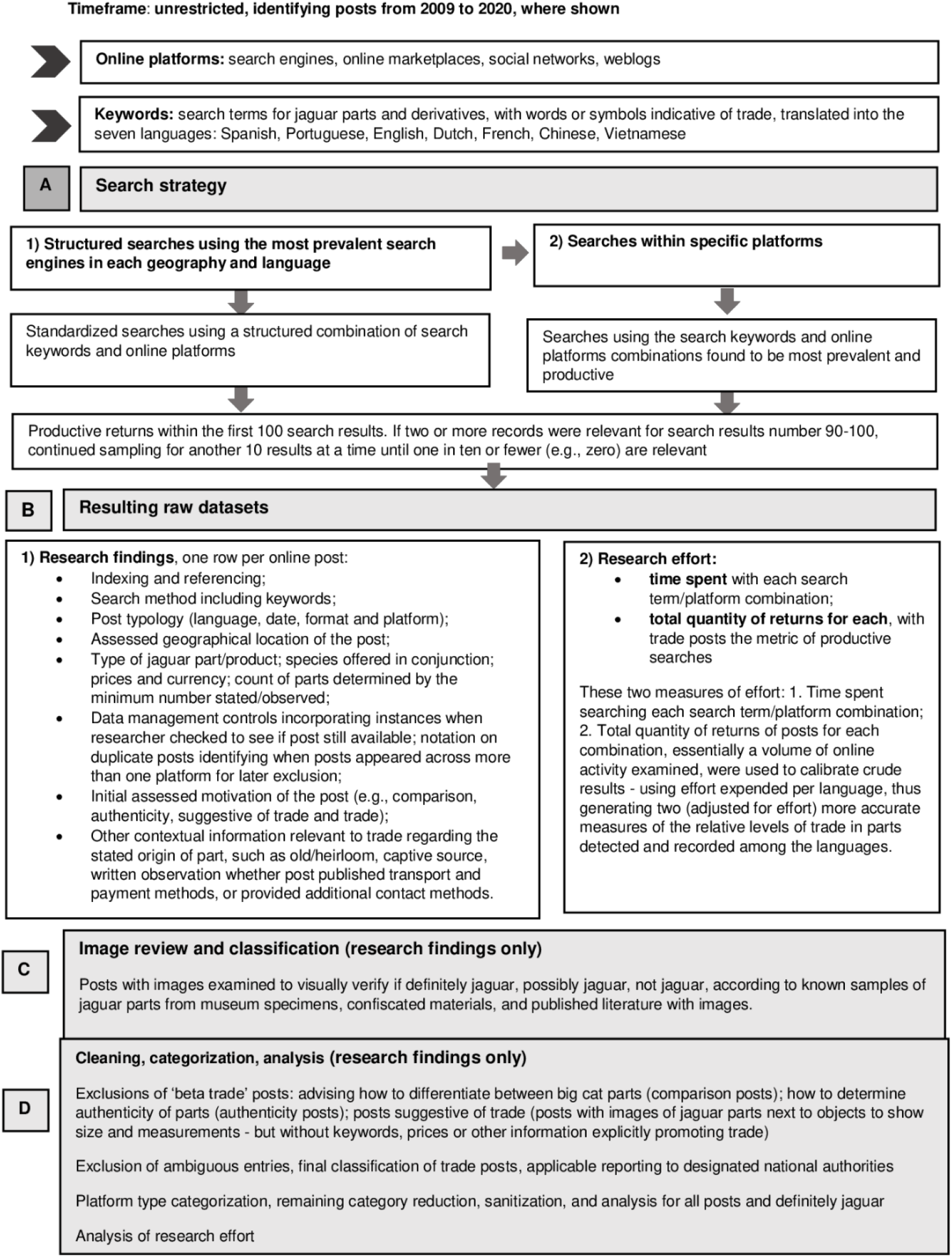
Flow Process Chart: Manual research of openly-available posts indicative of trade in jaguar parts and products.

### Ethics

Searches were conducted to generate replicable methodology for enforcement capacity-building efforts. Prior to commencing the project, all researchers undertook internal open-source research and ethics training and subsequently worked within defined research protocols addressing human rights, right to privacy, data protection, online research ethics, and risk mitigation in researching illegal trade.

We undertook manual searches to identify information placed in the public domain by internet users. We achieved this through using search engines which indexed search results, and searches within platforms incorporating search functionality. Whilst automated searches have been utilized in other studies [48,55,56], we considered them and noted that substantial pilot work using manual searches are a required preliminary step to design automated searches and machine-learning. Therefore, we used manual not automated methods, which whilst labor-intensive, enabled iterative development of search terms. Furthermore, some platforms restrict the use of automated tools, and we ensured consistency through deploying only manual searches.

Searches enabled us to view public posts, including those in open groups on platforms with such functionality, not private or ‘secret’ groups. We did not engage in entrapment or encourage illegal trade: we did not interact with prospective selling accounts, did not make bids on advertised parts, did not attempt to purchase any products, and did not verify physical presence or possession of alleged wildlife items. Validation of potential authenticity of advertised parts and products was therefore impossible, so we caution that results are indicative only.

We excluded search results pertaining to sharing of conservation news, along with display, wearing, or personal use of jaguar parts and products. We only analyzed posts assessed with confidence as motivated by and indicative of trade (Fig 1). This meant we assessed but excluded ‘beta trade’ search results which advised how to differentiate between jaguar and/or other big cat parts (comparison posts) or how to determine authenticity of parts (authenticity posts). Similarly, we excluded posts which implied but did not explicitly promote trade, for example posts with images of jaguar parts next to objects to show size and measurements - but without keywords, prices or other information explicitly promoting trade (suggestive posts), although we note such formats might be deployed to circumvent detection as trade. Our definition of trade therefore sought to reduce subjectivity in assessment, and it is likely that the results represent a conservative snapshot.

To ensure replicability and sharing of best practice methods for capacity-building, we restricted reviews of posts to publicly available information, reflecting law enforcement abilities prior to obtaining separate law enforcement-specific authorizations [69]. Furthermore, we did not create, merge, or integrate findings with additional datasets, practices which when using public data, can be considered invasive to privacy[70] even related to manual searches.

There are several means of reporting suspected violations online (see, for example, [71]). During the project, we followed legal obligations to report suspected violations directly to designated national authorities, and we provided the relevant URLs to designated government agencies, with appropriate handling and dissemination controls, for information purposes and for their further review. We note that for such specific purposes, URLs could not be anonymized. The project did not otherwise disclose or publish information which might identify specific posts, we have anonymized platform names, and do not publish prices or specific search terms.

## Results

Our team conducted 468 hours of searches, examining a total of 241,722 posts across the seven languages. We identified 230 trade posts with jaguar parts, of which 71 were accompanied by images identified as definitely jaguar, including a total of 125 jaguar parts. Using our conservative methods that screened out any ambiguous material this means we detected material from between 54 and 125 jaguars (95 teeth (24 to 95 individuals), plus 22 skins, 1 skin piece, 5 heads and 2 bodies). Spanish language searches found over a third of the total posts (91; 39.6%), and Chinese over a quarter (61; 26.5%) followed by Portuguese (44; 19.1%), Vietnamese (31; 13.5%), Dutch, English and French (1 each, 0.4% each). Following image review, just over half of definitely jaguar posts were found through Spanish searches (36; 50.7%), followed by Portuguese (18; 25.4%), Chinese (16; 22.5%) and French (1; 1.4%) (Fig 2). The search teams researching in Spanish did encounter some duplicate posts, which were therefore removed.

**Fig 2.**
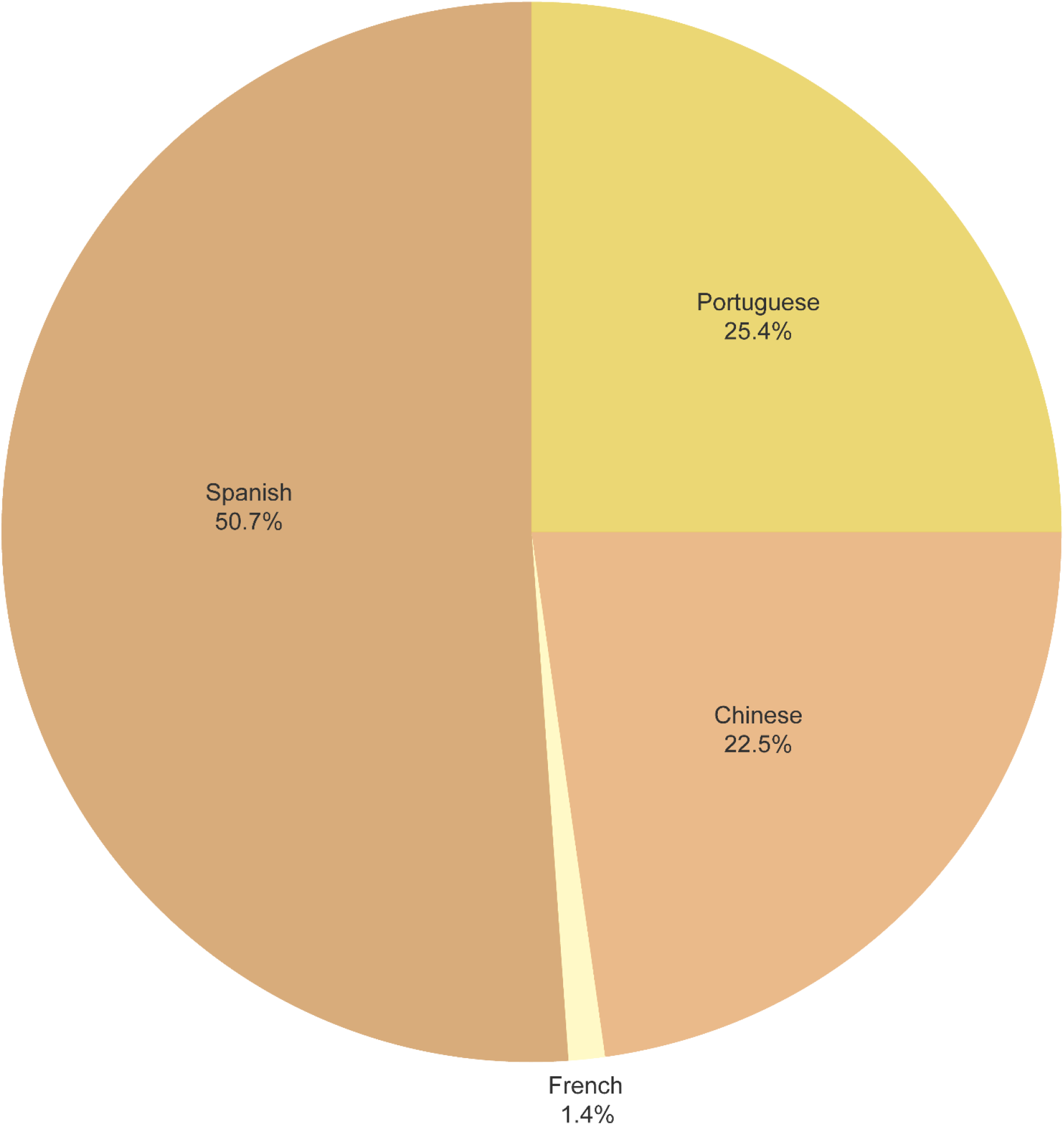
Percentage of posts identified as ‘definitely jaguar’ found in each search language (n=71).

The year 2018 was the most prolific year for identified posts (52 posts), but the definitely jaguar subset had a more even distribution across consecutive years from 2015 to 2019 (range 11-14 posts).

Where possible, researchers estimated the geographic location of the posts, through platform addresses, stated business geo-tag and other contextual information. We acknowledge the limitations in this approach and ‘locations’ are presented only related to assessment not verified. Posts were linked to at least 17 different countries; Mexico and Brazil had the largest shares of posts determined to refer to a country (Fig 3).

**Fig 3.**
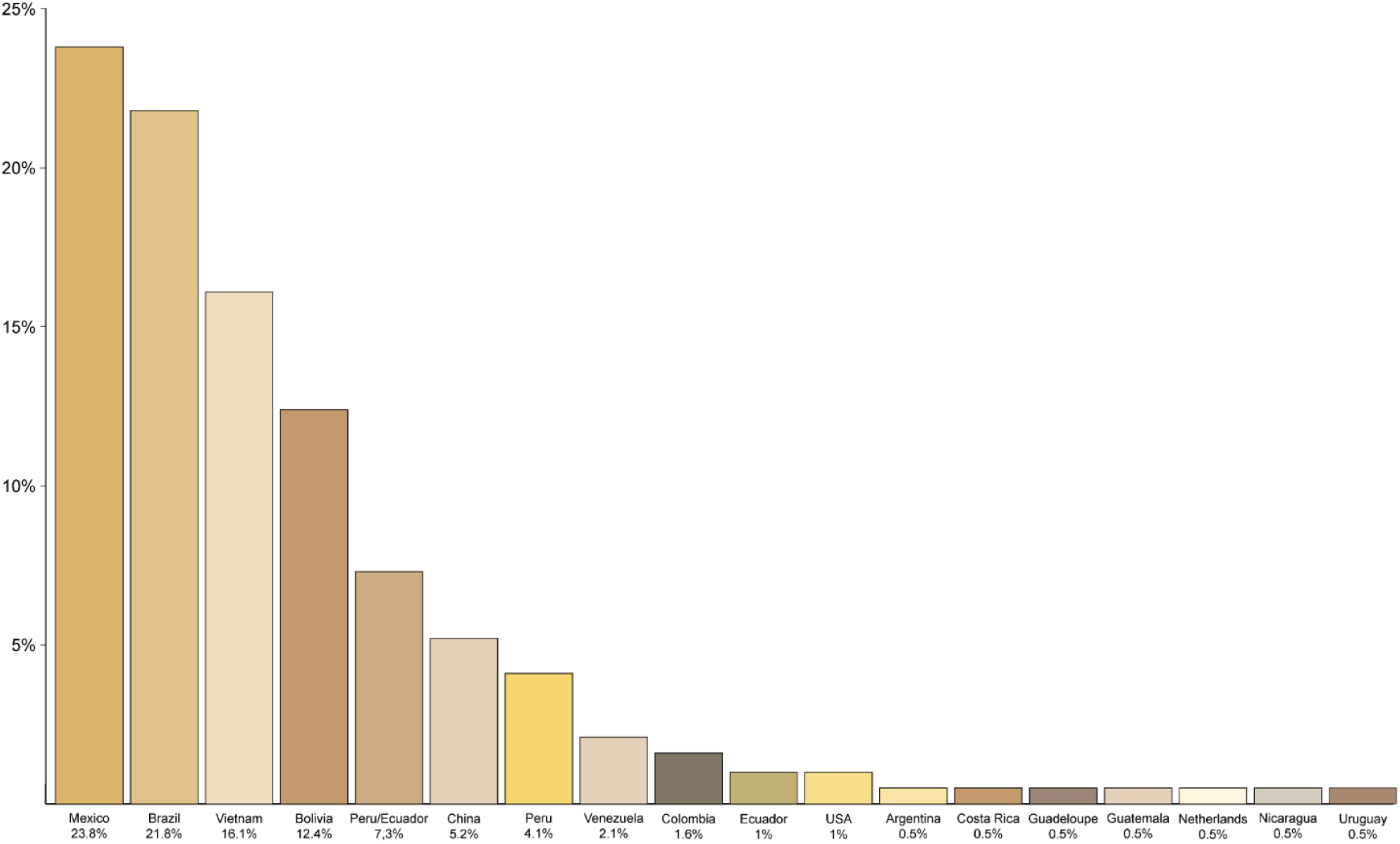
Percentage of posts advertising jaguar parts by country (n=193).

Posts were identified across 31 platforms. We found 230 posts across 21 online marketplaces (42 posts), 2 search engines (3 posts), 5 social networks (159 posts), 2 video-sharing sites (25 posts) and 1 weblog (1 post) (Table 1). On these platforms, 71 posts accompanied by images and assessed as jaguar appeared on 12 platforms, comprised of 10 online marketplaces (18 posts or 25.4% posts) and 2 social networks (53 posts or 74.6% posts) (Table 1, Fig 4). For the 125 jaguar parts, 93 parts (74.4%) were on social networks, and 32 parts (25.6%) on online marketplaces.

**Table 1.** Platforms by language (# of all posts with jaguar parts by platform and language; # of posts with visually confirmed jaguar parts, by platform and language)

| For each platform:<br>Row 1 = unverified<br>posts jaguar parts;<br>Row 2 posts<br>containing verified<br>jaguar parts | Chinese | Dutch | English | French | Portuguese | Spanish | Vietnamese | # Of languages by searching<br>platform type<br>Row 1, all posts with jaguar<br>parts;<br>Row 2, visually confirmed<br>jaguar parts |
| --- | --- | --- | --- | --- | --- | --- | --- | --- |
| Online<br>marketplace | 12<br>5 | 1<br>0 | 0<br>0 | 1<br>1 | 16<br>9 | 12<br>3 | 0<br>0 | 5<br>4 |
| Search Engine | 1<br>0 | 0<br>0 | 0<br>0 | 0<br>0 | 0<br>0 | 2<br>0 | 0<br>0 | 2<br>0 |
| Social Network | 48<br>11 | 0<br>0 | 1<br>0 | 0<br>0 | 28<br>9 | 76<br>33 | 6<br>0 | 5<br>3 |
| Video Sharing | 0<br>0 | 0<br>0 | 0<br>0 | 0<br>0 | 0<br>0 | 0<br>0 | 25<br>0 | 1<br>0 |
| Weblog | 0<br>0 | 0<br>0 | 0<br>0 | 0<br>0 | 0<br>0 | 1<br>0 | 0<br>0 | 1<br>0 |
| Count of posts by<br>language: Row 1,<br>all stated as jaguar;<br>Row 2, visually<br>confirmed as<br>jaguar | 61<br>16 | 1<br>0 | 1<br>0 | 1<br>1 | 44<br>18 | 91<br>36 | 31<br>0 | NA<br>NA |

**Fig 4.**
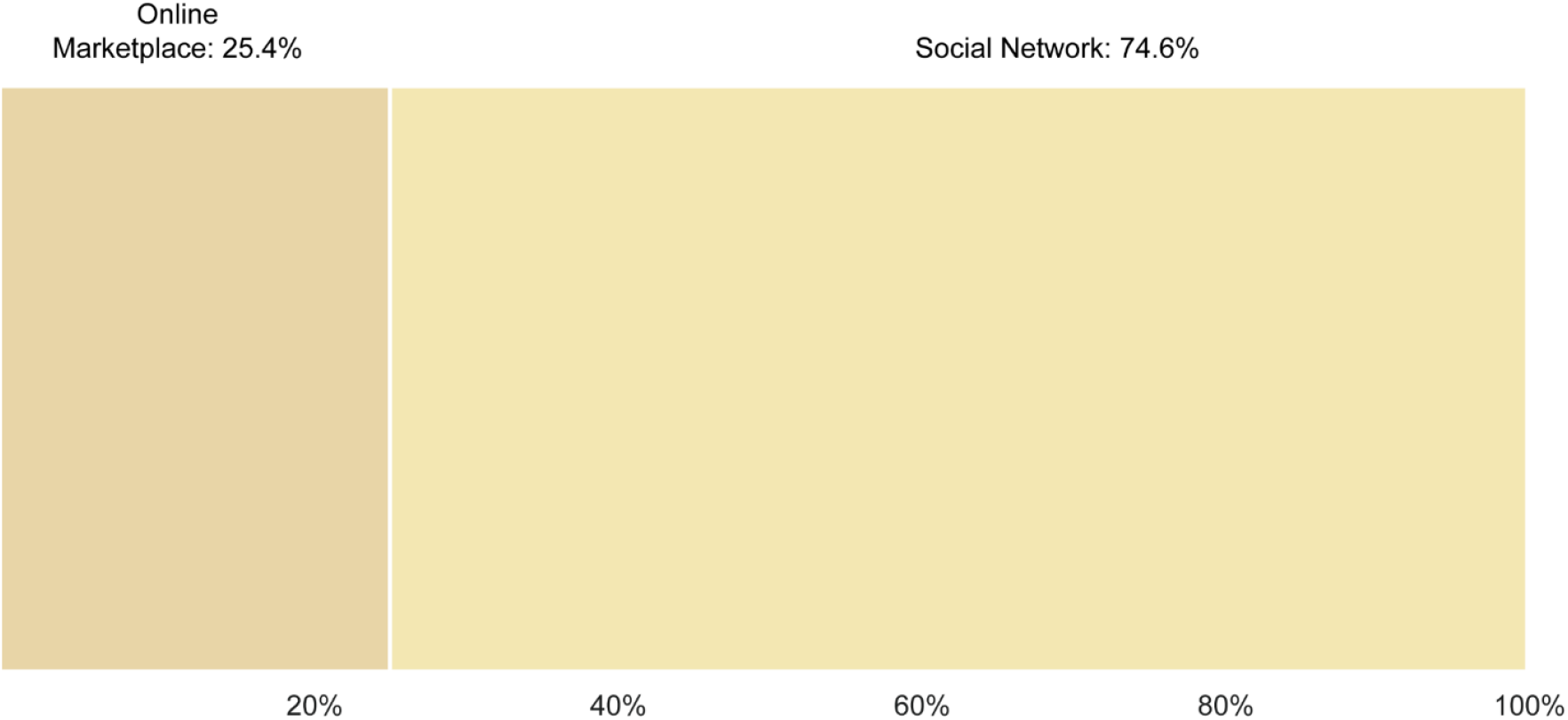
Percentage of visually verified jaguar posts on social networks and online market places (n=71).

All platforms were searched in at least one language, one platform in 2 languages and one platform in 5 languages. The presence of more than one language on some platforms resulted in 11 platforms showing posts in Portuguese, 11 platforms in Chinese, 7 platforms in Spanish, 4 platforms in Vietnamese, and 1 platform each in Dutch, English and French. Spanish searches found the greatest number of posts assessed as jaguar (36 posts located on 2 platforms), whereas Portuguese searches found fewer jaguar posts however across a greater number of platforms (18 posts located on 6 platforms) (Table 1).

Parts were identified across 15 different categories, 14 harmonized with the UNEP-WCMC CITES trade terms (https://trade.cites.org/) plus whiskers, and those which were definitely jaguar were bodies, heads, skins, skin pieces, and teeth.

We found that teeth were the most prolifically traded part online; 156 posts offered at least 367 teeth; 42 of those posts featured at least 95 teeth, as featured in images and assessed as definitely jaguar (Fig 5).

**Fig 5.**
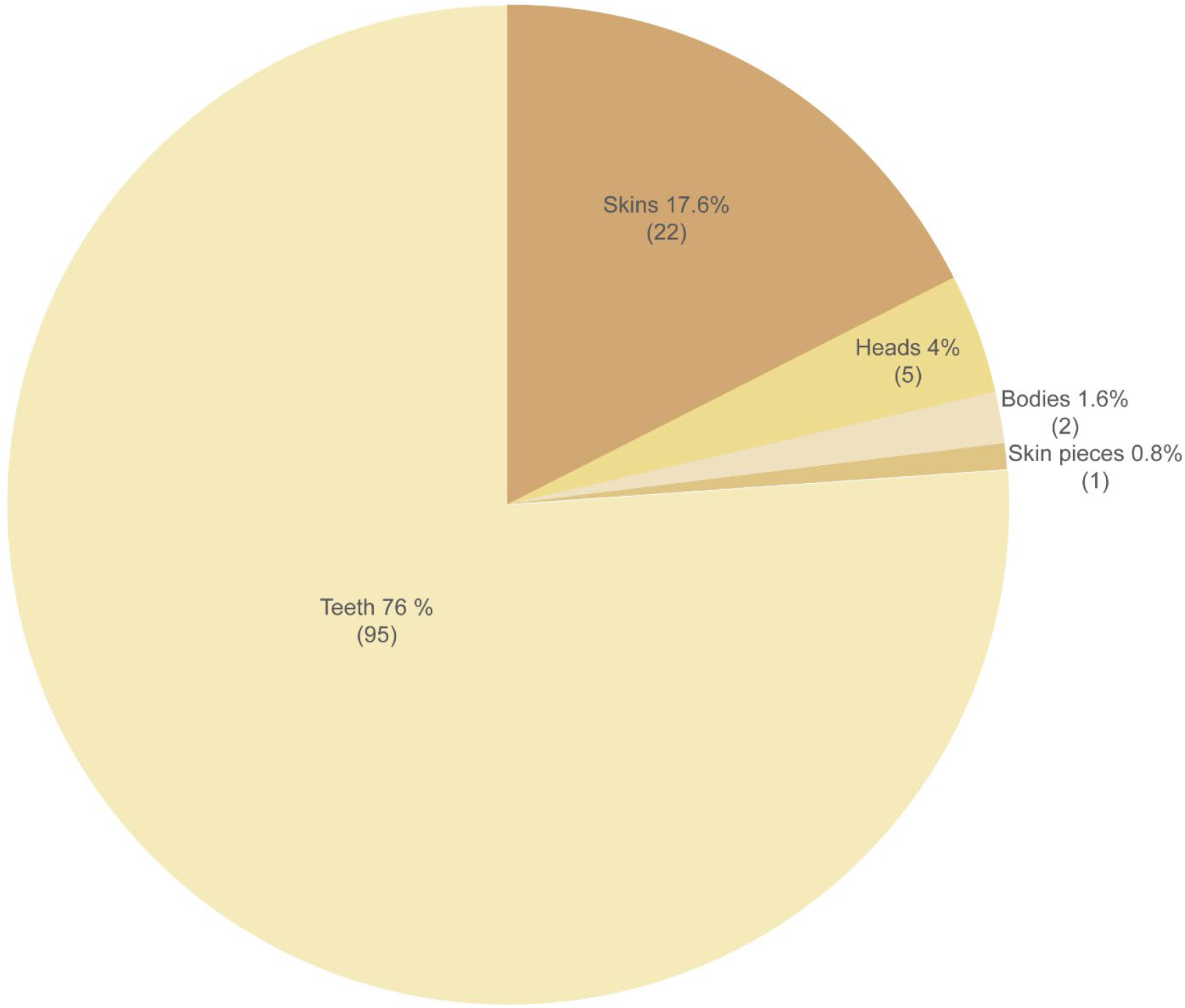
Number and percentage of confirmed jaguar parts by type (n = 125).

Of these, 71 teeth were assessed as jaguar and linked to a location (Fig 6a, within which Mexico and China had roughly one quarter each of these teeth (26.8% and 25.4% respectively), followed by Bolivia and Brazil (16.9% and 12.7% respectively). Without adjusting for effort, the number of teeth by country/dual country was Mexico (19), China (18), Bolivia (12), Brazil (9), Peru/Ecuador (most accurate probable location) (8), Venezuela (3), Guadeloupe (1), and Uruguay (1).

**Fig 6.**
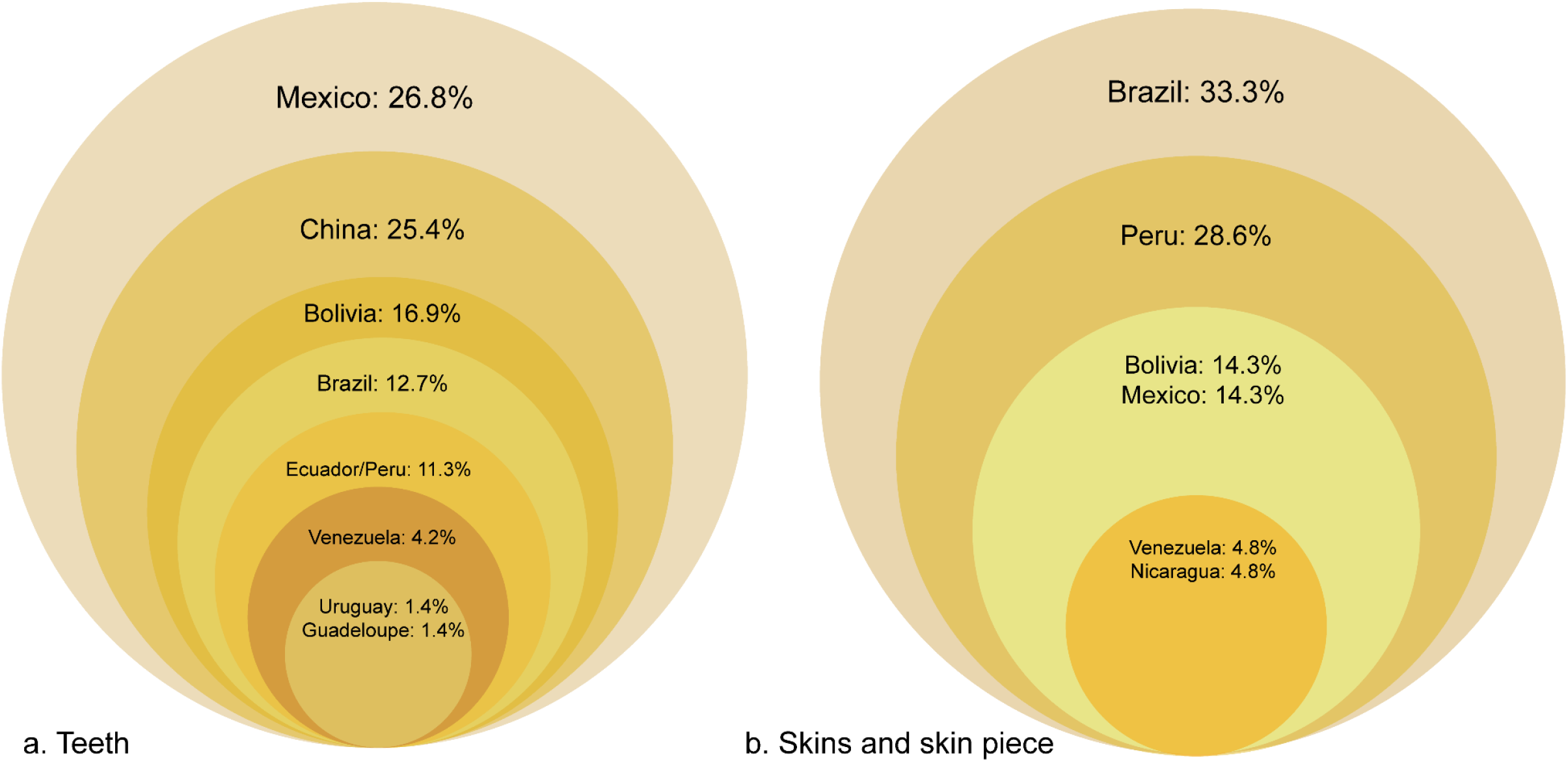
Fig 6a. Percentage of detected jaguar teeth linked to a location in each country (n=71). Fig 6b. Percentage of detected jaguar skins and skin pieces linked to a location in each country (n=21).

Skins comprised the second most-traded part; 37 skins with additional skin pieces and scraps featured (including in posts showing more than part), and 22 skins with 1 additional skin piece were definitely jaguar. When an assessment of location of post was possible, skins were identified only in Latin America, therefore from that subset, 20 skins were assessed to link to a location: Brazil (7 skins = 35.0%), Peru (6 = 30.0%), Bolivia (3 = 15 %), and Mexico (2 = 10%), followed by Venezuela and Nicaragua (1 skin each = 5.0% each; all referring to Species 1 column only).

If the additional skin piece is also considered, (n=21), these are Brazil (7 skins = 33.3%), Peru (6 skins = 28.6%), Bolivia (3 skins = 14.3%), and Mexico (2 skins plus 1 skin piece, n=3, = 14.3%), Venezuela and Nicaragua (1 skin each = 4.8%) (Fig 6b).

We identified 12 posts totaling at least 14 claws. Although we could not confirm that they were jaguar based on images, claws were connected to Mexico (sum of 12 claws) and Costa Rica (1).

The body parts we encountered as ranked by number of image-verified posts and referring to locations were teeth (36 posts), skins (20, plus 1 post for skin piece), heads (5), bodies (2) (Table 2), and by sum number of parts as teeth (71), skins (20 plus 1 skin piece), heads (5) and bodies (2) (Table 2). A small number of bone posts were identified through searches, but not verified by images. Research gained insufficient evidence to assess the prevalence of online bone trade in countries.

**Table 2.** Visually confirmed jaguar body parts by country or region (# posts per country or region of posting; sum of parts by country or region of posting)

| Row 1 of each country = # Posts<br>Row 2 of each country = # Parts |  | **Bodies** | **Heads** | **Skins** | **Skin pieces** | **Teeth** | **Total** |
| --- | --- | --- | --- | --- | --- | --- | --- |
| Bolivia | Number posts |  | 4 | 3 |  | 4 | 11 |
|  | Sum of parts |  | 4 | 3 |  | 12 | 19 |
| Brazil | Number posts | 2 |  | 7 |  | 9 | 18 |
|  | Sum of parts | 2 |  | 7 |  | 9 | 18 |
| China | Number posts |  |  |  |  | 4 | 4 |
|  | Sum of parts |  |  |  |  | 18 | 18 |
| Guadeloupe | Number posts |  |  |  |  | 1 | 1 |
|  | Sum of parts |  |  |  |  | 1 | 1 |
| Mexico | Number posts |  |  | 2 | 1 | 11 | 14 |
|  | Sum of parts |  |  | 2 | 1 | 19 | 22 |
| Nicaragua | Number posts |  |  | 1 |  |  | 1 |
|  | Sum of parts |  |  | 1 |  |  | 1 |
| Peru | Number posts |  |  | 6 |  |  | 6 |
|  | Sum of parts |  |  | 6 |  |  | 6 |
| Peru/Ecuador [nearest locational assessment] | Number posts |  |  |  |  | 5 | 5 |
|  | Sum of parts |  |  |  |  | 8 | 8 |
| Uruguay | Number posts |  |  |  |  | 1 | 1 |
|  | Sum of parts |  |  |  |  | 1 | 1 |
| Venezuela | Number posts |  | 1 | 1 |  | 1 | 3 |
|  | Sum of parts |  | 1 | 1 |  | 3 | 5 |
| Totals | **Number posts** | **2** | **5** | **20** | **1** | **36** | **64** |
|  | **Sum of parts** | **2** | **5** | **20** | **1** | **71** | **99** |

Without adjusting for effort, an abbreviated ranking of countries/regions from which we identified posts with images of verified jaguar parts is as follows: Brazil (18 posts), Mexico (14), Bolivia (11), Peru (6), Peru/Ecuador (most accurate probable location) (5), and China (4), followed by further inclusions (Table 2). Effort was variable. Simply looking at pre-data filtering effort, the time spent searching ranked by minutes was Spanish (21,046), Portuguese (2,590), Vietnamese (1,859), Chinese (1,090), English (779), French (546), and Dutch (190). In total, the team searched for 468 hours, 20 minutes, and 41 seconds. However, the number of “search term/platform combinations returns total” as an indication of how many online searches (unique search term/platform combinations) actually transpired, whether yielding “productive returns” or not pre-filtering, contrasted; Spanish (203,783 returns); Chinese (14,950), Portuguese (11,944), Dutch (5,529), French (3,349), English (1,414), and Vietnamese (753). Note how four times as much searching was spent in English than Dutch, however, in that time, there were nearly four times as many searches in Dutch, indicating that the scrutiny given to each return was another source of sampling variation. Vietnamese ranked third in time spent searching, and seventh in searches completed. The returns/effort can also be expressed as relative frequencies (time per language/time total and searches per language/searches total, with both summing to 1.00), but are more easily visualized as numbers (of minutes and searches).

Considering that we had two orders of magnitude of variation in time spent per language, and three orders of variation in actual searches, our crude results have considerable sampling bias; highlighting that effort was not consistent across languages and geographies. However, we can assess an adjusted “hit rate” (productive results by time and searches). This “equalizes” variation across languages and geographies. The ranked “relative frequencies of productive returns” (productive search returns as a proportion of all returns by language, before data filtering and image verification) were Vietnamese (0.0996), Portuguese (0.0148), English (0.0042), Chinese (0.0039), Spanish (0.0019), and Dutch (0.0002). Overall, relative frequency of productive searches per searches total was 0.0029 or 0.29% of all 241,722 searches. When divided by time, the rank changes, with productive returns per total search time in minutes was as follows: Portuguese (0.0683), Chinese (0.0541), Vietnamese (0.0403), Spanish (0.018), French (0.0092), English (0.0077), and Dutch (0.0053). Overall relative frequency of productive searches by time was 0.025 or 2.5% of each of the 28,100 minutes invested.

The above ranks are before image classification and data filtering. As mentioned above, because of ambiguities of language, upon closer examination, many Vietnamese posts were identified as leopards, bears, tigers, and unknown, but none as definitely jaguar. Removing the false positives in Vietnamese and other languages, the results in a ranked relative frequency series were Portuguese (0.0148), English (0.0042), Chinese (0.0039), Spanish (0.0019), French (0.00015), and Dutch (0.0002) with 0.26% of the 240,969 search returns being positive, and ranked relative frequencies of Portuguese (0.0683), Chinese (0.0541), Spanish (0.018), French (0.0092), English (0.0077), and Dutch (0.0053) by time with 2.39% of the searches per minute (26,421 minutes total) positive.

There are contrasts in country area and population sizes, and thus in the use of languages within the jaguar range. For example, French Guiana has a population of 299,000 people in 82,000km², Suriname has 587,000 people in 156,000km², Belize has 388,000 people in 22,810km², and Guyana has 787,000 people in 196,850km², totaling 1,175,000 people in 219,690km², whereas Brazil contains 213,000,000 people in 8,358,140km² [72]. The varying scales that each language is employed across the arena of online jaguar trade may be reflected in research results.

Given that search effort varied by orders of magnitude among languages, we cannot responsibly adjust the crude results using proportions of effort. However, the relative proportions of where we encountered online trade in jaguar parts did change when adjusted for effort. Since these are relative frequencies, the country’s size plays no factor in the following ranks. Standardizing by dividing by effort expressed by search time and also by number of individual searches using search term/platform combinations (and excluding Vietnamese), the ranked prevalence of posts with jaguar material were Portuguese (Brazil) #1 by both metrics. Chinese #2 and #3 (per time, and per number of searches, respectively), Spanish ranked #3 and #4, mainly due to Mexico, Peru, and Bolivia. English ranked #5 and #2, and Dutch ranked last in our searches. These geographical proportions, based on data from a semi-systematic investigation of online trade, contrasted with reports focused on individual countries, and from some aspects of the CITES report that primarily relied upon an aggregate of official records (including seizures), country specific primary research, and literature searches [9].

By recording search term/search engine/online platform combinations and sorting by proportions of productive returns, we were able to identify the most productive combinations. We decline to publish these terms due to unknown potential impacts upon online trading behaviors.

Despite all the caveats that we have presented, the methodology we developed increased our understanding of the widespread trade in jaguar parts, and advancing approaches applicable to other taxa. These methods have since been used on frogs, turtles, crocodilians, boas, parrots, macaws, painted buntings, ocelots, pumas, peccaries, and primates.

## Discussion

Our research presents a snapshot of prevalence of online jaguar trade, and serves as a contribution to current and ongoing efforts to better understand the threats to jaguars in the wild. In the process, we tested methods of conducting investigations of online trade that have utility for many species now traded online. We found evidence of online trade within jaguar range countries, and in Asia, whilst revealing knowledge gaps which may be addressed through further research. We found that jaguar teeth are the most prolifically traded parts online. Interpretation of that has to factor that each jaguar has four canines. Despite keyword searches retrieving posts in Vietnamese, our subsequent image verification process was unable to identify any jaguar teeth in Vietnamese posts. Online trade in tiger parts has been identified related to Viet Nam [62] and image assessment suggested tiger or other big cat parts traded rather than jaguar.

In contrast to Vietnamese language posts, Chinese language posts had a higher proportion of posts advertising jaguar teeth that image review classified as genuine. Furthermore, we observed some ‘beta trade’ Chinese language search results (not analyzed) discussing how to determine authenticity and species differentiation, suggesting a potential market for jaguar teeth. However, further research is needed to estimate the contemporary prevalence of any domestic market in China for jaguar parts and, as relevant, determine drivers of consumption. Our research did not find jaguar skins advertised in Chinese and assessed as within China, and no skins advertised in Vietnamese; therefore, jaguar skin trade there or other Asian countries remains unconfirmed and warrants further research.

Notably, despite several reports of a paste or glue derived from boiled jaguar bones in Suriname that is sold within that country and linked to Chinese communities [16,38,40], our survey found scant credible posts of bones, with no verifiable images, whether in Dutch, Chinese, or Spanish, and no posts of paste. This does not negate the possibility that the practice and trade exist, only that these methods and this survey did not find noteworthy evidence. We are not aware of identified jaguar bone paste seizures in China.

## Trading exploitation of multiple platforms and species

Nearly one fifth of posts (44 posts or 19%), particularly on video-sharing platforms (23 posts) published other details such as trading address, trading websites, email addresses, and additional platform contact methods or channels. We did not analyze these, but we consider that law enforcement investigations of online trade might include concurrent exploitation of multiple platforms as one element of threat and prioritization processes.

The Wildlife Justice Commission (WJC), which investigates illegal wildlife trade, found some users may openly advertise available stock on one platform, but negotiate business through closed messaging platforms [27], echoed by [73]. The United Nations Office on Drugs and Crime notes that middlemen and sellers use private groups to reach customers and switch between platforms [29]. Our study did not address private channels or groups and therefore an unknown proportion of trade may take place through such means.

Few posts published specific instructions on payment methods, with 16 posts instructing to contact the seller, 6 suggesting transfer service or payment platform, and 1 further post stating cash, credit or debit card. Few posts published an offered transportation method, but those that did mostly indicated postal service (n=8) or courier (n=1), with even fewer posts referring to personal collection or to be agreed.

Only 4 posts concurrently offered more than one jaguar part (recorded in ‘Species 2’ column). Therefore, whilst we have calculated the sum of parts to include co-occurring parts, only 2 of the posts were assessed as definitely jaguar, but that assessment refers to the primary-listed parts only (‘Species 1’ column). In some instances (23 posts, or 10% of the 230 posts) posts co-occurred with other species, including as claimed and not assessed: tiger, puma, crocodile, peccary, sea lion and shark; 3 posts with genuine jaguar images also showed crocodilians and puma.

Researchers reviewed posts to examine if parts were specified as captive source, however only 1 post suggested this but assessed as not jaguar. In addition, few of all posts stated the parts were old (n=5) and inherited (n=1). Due to lack of information, we were unable to further analyze the potential contribution of either alleged captive-sourced or old parts to trade, but recommend these factors are taken into account in future studies.

Encouragingly, whilst Chinese language research retrieved posts within search results, attempts to view some posts were blocked by platform screening processes, which shows the contribution which platforms can make to limiting content.

## Geographical variation in trade

Most reports of illegal trade in jaguar body parts prior to this study focused on Bolivia, Suriname, and China [14,16,38,40]. Where we could assess ‘location’ of posts, we found Mexico was linked to a greater number of teeth than China and Bolivia, and Brazil and Peru had more posts of skins than Bolivia, although all were low in numbers (Table 2). These findings present a snapshot of online trade which is not to diminish previously-identified threats whether online or offline. However, they do emphasize that jaguar trade has a greater geographical scope than it appeared based on coverage of the issue prior to 2021. The greater volumes of material that we encountered in Brazil and Mexico, in contrast to Bolivia and Peru is notable, as is that we detected trade emanating from China. Even when we standardized for effort and before further triage, Portuguese and Chinese remained leading languages as far as number or productive posts. The geographical proportions of trade that we encountered were not a tight match with the summarized seizure data presented in the 2021 CITES report [9] but the explanation is probably simple. Some (though not all) of the data in the CITES report emanated from global seizure records and official national reports. Although the CITES report [9] included primary research, literature research, expert input, and other sources (including the preliminary results of this investigation), the seizure record portions may have been positively biased towards the countries with the most efficient wildlife trade enforcement (e.g., USA and Germany). Conversely, official poaching and trade data based upon interceptions in many jaguar range states have been relatively thin, introducing a potential negative bias; where trade exists but is under-detected and/or under-reported. We designed a relatively comprehensive semi-systematic sampling of online trade, with an effort to standardize results across languages. Take-home messages of the CITES report [9], which this study does not weaken, are that; 1) across jaguar range, improved data collection and storage is urgently needed; 2) a wide range of sources needs to be considered for a comprehensive understanding of the illegal trade in jaguar parts. The specific contributions of this study are: 1) pilot methods for conducting online investigations to understand and counteract IWT in jaguars; 2) the results of a pilot investigation across a ten-year band and seven languages.

## Internet availability and utilization

Internet access and use varies across countries, and within national populations (S5 File Table). As a consequence, the degree by which commerce is conducted on online platforms varies among countries [51]. The proportion of populations in our study area with access to the internet varied from 93% to 37%, with Argentina the country with the highest access to internet and Honduras the lowest [74]. Brazil has an above average internet penetration for the region (70% of the population), which, combined with a large area of remaining habitat, may explain its large number of reports (Fig 7). Mexico, Bolivia and Peru present a similar proportion of the population with internet access (∼67%) [74].

**Fig 7.**
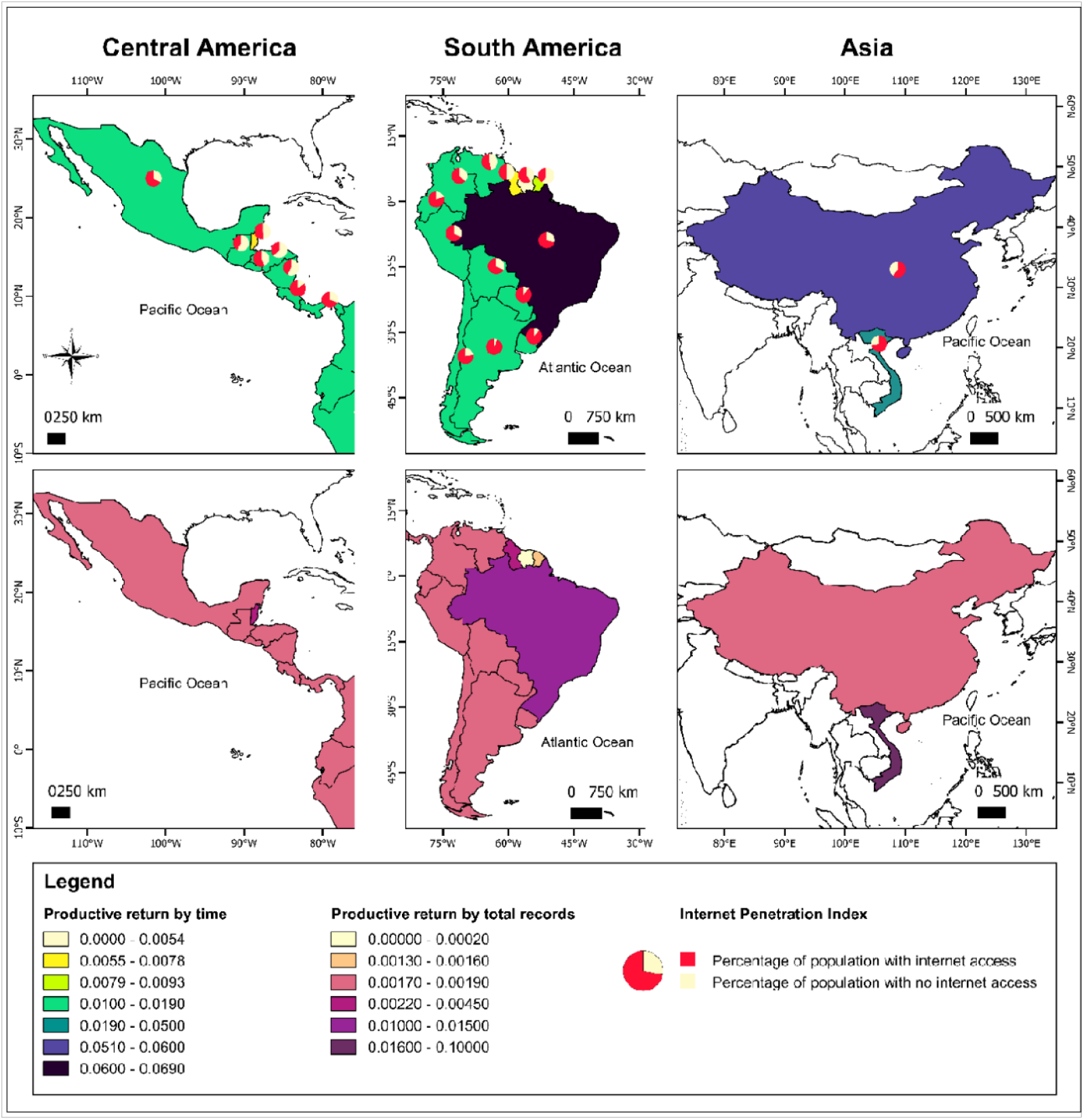
Internet penetration index and returns. Internet penetration index [74] and returns (by language and country) related to productive returns by 1) time spent searching; and 2) total records (searches executed). This is prior to verification of images as jaguars and thus is more reflective of the volume of online trade in felid and other carnivore parts than specific trade in jaguar parts.

Overall, we identified less evidence from the Guianas (Guyana (English), Suriname (Dutch), French Guiana (French)) than expected, especially in the context of previous reports based upon physical surveys and questionnaires [16,38,40]. However, we lack a confident explanation to attest whether trade is rarely occurring in those countries, or is not prevalent online, or takes place on platforms not accessed. Suriname, Guyana, and French Guiana (60%, 50% and 41% of the population with access to the internet) experience an internet penetration up to 36% lower than the average for Central and South America [74]. In remote frontier areas, electricity is unreliable, for example over 90% of Suriname is forested, which would make online trade challenging.

Human population size is also unequal among countries, as is the number of people accessing the internet. Cayenne, the capital of French Guiana, has a population of 61,550 [72]. In comparison, Iquitos, Peru, has a population of 437,620, Lima, Peru, 7,737,000, and Mexico City 12,294,193 [72]. Given their lower populations, and substantial portions of remote forested areas, Suriname, Guyana, and French Guiana together represent the equivalent of 0.2% of total internet users of Central and South America [74]. Conversely, Brazil and Mexico, the two top countries in number of records, together hold more than half of all internet users (57%) in the sampled region [74]. Therefore, online investigations in jaguar trade carry inherent sampling bias and may be improved by complementing the online search with physical observations, for example, from market surveys and seizures.

However, even in countries where internet penetration is on average high, such as Brazil, access to the internet may vary considerably among regions within the country. The percentage of households with access to internet is higher in urbanized federative units, while remote places closer to jaguar habitat presents, in general, lower internet access. The importance of internet as a trading medium may vary correspondingly across these gradients (S5 File Internet Penetration).

## Sampling challenges and bias

Defining search terms is a challenge in online wildlife trade, which is constantly evolving. Post content may avoid using specific terms indicating trade and may employ slang or emojis, and there are vernacular names in general use for jaguar in different countries and regions which may potentially be used, including the use of some indigenous terms mixed with terms in the official country language. Xu et al. [73] recommend robust surveillance in multiple languages, incorporating localized keywords and vocabulary. We did exactly that by constructing an extensive list of words used in each language/country, but agree that the language and symbols that traders use will evolve, requiring some agility to track.

Chinese and Vietnamese language characters do not differentiate between different big cat species to the extent of some other languages. Mandarin Chinese does not have a specific word for jaguars [75] and the term American leopard (美洲豹) is used (or American tiger, 美洲虎, which is less used). In such cases, contextual trade terminology helped to identify postings, images from which were reviewed for authenticity. For example, contextual additional keywords included the equivalent of ‘the Americas’ and ‘South America’. Since sampling variation (and bias) is a risk in any investigation, a carefully considered design for online research of wildlife trade with as much cross-language and geography standardization is recommended. Sampling constant proportions of a ‘population of interest’ will result in robust indices [66]. When sampling proportions are not constant, raw results may be as much a metric of variation in search effort, search terms, and search types as a measure of the ‘population of interest’. Even with standardized sampling protocols, the search terms, platforms, and time and search effort (returns) employed should be carefully recorded to aid interpretation.

## Conclusions

We found evidence of openly-available online illegal jaguar commerce in several countries. These findings expanded previous perceptions of geographic patterns prior to this study, with high proportions of validated detections coming from Brazil and Mexico. Although the scale of material (between 54-125 jaguars) may seem low given the ten-year sample band, it is important to consider the following: 1) the estimates are conservative; our visual methods for inclusion rejected all material of ambiguous identity (including, but not limited to, images too poor to definitively assign to jaguars); 2) this represents the proportion of all online trade in jaguar parts that we detected, not necessarily all that was traded; postings could have been taken offline following sale or as detected by platforms; 3) an unknown proportion of trade may occur on additional platforms and in groups with controlled access, which we intentionally did not access for ethical reasons; 4) as in any wildlife research, detection probabilities are rarely 100%. The findings complement other research methods to refine our understanding of where online trade is most prevalent. As an example, a subset of this sampling deployed the exact same research effort in Paraguay and Argentina as Bolivia. In that sub-sample, posts of jaguar parts were detected and recorded from Bolivia, but not the other two.

Our findings confirmed that online trade in jaguar parts occurs, and may serve as a baseline to track future trends. An inference we draw from this research and other recent studies [9,15,18,19,20] is that even after the jaguar being listed on CITES Appendix I drastically reduced international trade, low-grade yet common use of and trade in jaguar parts persisted in some range countries/regions. Recent concern about re-emerging international trade [14,16,21,38], that with links with Asia and/or Asian diaspora [9,19], drove a surge of research [15,18,20] that has revealed the extent of persistent local and national trade, and that can help jaguar range countries address the full spectrum. The widespread extent of commerce we uncovered emphasizes the need to directly curb all three levels of trade: local, national and international. There are implications for policy and enforcement including the need for increased vigilance and additional investigations to better understand and devise appropriate responses to curb this threat. Future research should include enhanced understanding of demand drivers, supply chains, collection and transportation methods. Improved capabilities and commitments towards prosecuting poaching and trafficking cases is also necessary. The need for improved implementation of existing legislation is clear [76]. Demand reduction efforts should take place both specific to jaguar and within the context of other big cat trade, as relevant to different markets and geographies.

Existing partnerships between researchers, enforcement, and the technology sector can benefit from enhanced knowledge of the methods and evolution of online jaguar trade. We recommend consistent monitoring of both physical and online markets to increase understanding and identify emerging patterns and threats, complemented by governments maximizing mechanisms and social awareness programs to disrupt and discourage trade in jaguar parts at local, national, and international levels - law enforcement can be complemented by social awareness programs that reduce the social acceptability of trade in jaguar parts and communicate the penalties. However, those penalties need to be appropriately exacted to serve as a deterrent.

To some degree our findings across many countries echo previous studies in Belize and Guatemala [18], with considerable domestic/local traffic with weak national institutional responses appearing to allow an almost unregulated trade. Poorly managed human-jaguar conflict, outdated legal systems, inadequate field resources, staff, capabilities and efficient legal processes, and systematic corruption, weaken the ability of existing laws to temper jaguar trade and these drivers need to be addressed to curb the threat [18,19].

We note that all Latin America jaguar range countries have prohibited trading of endangered species parts and products, and people found trading jaguar parts and products are liable to administrative and criminal sanctions [76]. Adequate enforcement of these regulations will determine whether wildlife parts and by products end up being traded after being allegedly hunted for reasons other than trade. No trade of jaguar parts should be legal regardless of origin.

Arias et al. [18] found that opportunistic hunting tied to human-jaguar conflict was a source of trade in jaguar parts in Belize and Guatemala. Methods to reduce jaguar-livestock conflict have been developed to minimize jaguar mortality, and thus the potential for trade [67,77,78,79,80]. We recommend using conflict reduction protocols to minimize jaguar mortality (and parts thus entering trade channels), but we must emphasize that directly curbing trade can eliminate financial incentives to kill jaguars for parts, thus facilitating efforts at human-jaguar coexistence [15] and improving jaguar survivorship.

A combination of policies, prevention and deterrence mechanisms, and reinforced implementation efforts is needed to stop illegal trade. Since legislation in each country regulates against trafficking in the parts of threatened and endangered species, including jaguar, there exist opportunities and a strong need to deliver jaguar protection more actively and effectively in both rural areas where the killing happens and urban areas where demand for parts may exist. Studies, such as this one, seek to inform such efforts.

Immediately following this research, strict border and travel restrictions raised during the early months of the global COVID 19 pandemic reduced IWT [81]. Morcatty et al. [52] noted that pandemic caused delivery delays and discounted prices in general online wildlife trade in Brazil and Indonesia. However, concern about spreading zoonoses barely entered the conversations of practitioners and the authors found no clear evidence that the volume of online wildlife trade decreased. Conversely, reduced field patrols and diversion of law enforcement resources to COVID-19 issues had the potential to allow an increase in poaching and protected area invasions [81]. That potential appears to have been realized in a parts of jaguar range [82,83]. In addition, there is a possibility that illegal online trade in wildlife (“working from home”) flourished during the pandemic [84].

The methodology developed in this investigation has been modified for multiple taxa in the Andes-Amazon region, and, with refinements, has since been deployed in Mexico. Thus far, these methods, and refinements on the foundation they provided have been used on frogs, turtles, crocodilians, boas, parrots, macaws, painted buntings, ocelots, pumas, peccaries, and primates. We offer the general design and our findings in hopes of advancing efforts to combat IWT.

In August of 2019, the CITES Secretariat issued a decision to commission a study on the illegal trade in jaguars [85 CITES Decision 18.251, paragraph a], a study now complete [9]. In October 2019, the first High-Level Conference of the Americas on Illegal Trade in Wildlife recommended that IWT conducted through the internet be combatted with effective sanctions and penalties [1]. In February 2020, the jaguar was listed in Appendix I and II of the Convention for Migratory Species [86]. The methods and findings in this study can facilitate these decisions, and support the 2030 Jaguar Conservation Initiative [4] to make advancements in saving the great cat of the Americas.

## Supporting information

S1 - Condensed project plan

S2 - Dataset used for results analysis

S3 - Effort log template

S4 - Guidance - Standardization and identification of efficient search term & platform combinations

S5 - Internet penetration

## Acknowledgements

We thank the US Fish and Wildlife Service for support during this study, as well as the support of the European Union to address wildlife trade in the Amazonian countries, and the U.S. Department of State Bureau of International Narcotics and Law Enforcement Affairs (INL) to address these issues more broadly across Latin America. The Wildlife Conservation Society (WCS) partially supported most authors during this period and Melissa Arias and Thais Morcatty were also supported by WCS Scholarships. We thank Dr. David Roberts for advice given while developing our study designs, while organizing our data, and for a review of the manuscript draft. We are grateful to Guido Miranda and Omar Torrico for assistance in production of the figures. We dedicate this report to Julio Alfredo Madrid Montenegro, for the depth of his contributions to conservation, and in particular, his role in advancing this project’s goals even as COVID 19 cut his life short in September of 2020.

---

S1 File: Condensed project plan

S2 File: Dataset used for results analysis S3 File: Effort log template

S4 File: Guidance standardization and identification of most efficient search term-platform combinations S5 Table: Internet penetration, number of internet users, and percent of internet users per country

