## Supplementary material for "Multi-lingual multi-platform investigations of online trade in jaguar parts": S1 - Condensed project plan

### OBJECTIVE AND ACTIVITIES

We proposed an action-oriented summarization of spatial and temporal trends in trade in jaguar parts. We comprehensively gathered and systematized all accessible available information on jaguar trade through proactive monitoring of online platforms in multiple languages. Our searches were conducted in seven languages: Chinese; Vietnamese; English; Spanish; Portuguese; French; Dutch. This contained three levels: 1) systematic searches on diverse online platforms using standardized sampling, search methods and data collection methods, with search terms translated in various languages; 2) a second round of searches using the search term and platform combinations found to be most prevalent and productive for jaguar trade using standardised sampling, search methods and data collection, with search terms translated in various languages; 3) later in the project, for deeper examination, dive down searches in known platforms, without constraints of standardization.

We understood number 1 and 2 above could inform number 3, although number 3 could also be informed by professional knowledge of potentially productive platforms. In addition, the first phase of number 1, using multiple search terms and online platforms, informed the more refined and time efficient second phase nested within the standardized component. Such revisions were judged and made at mid-term mark in our study.

### SPECIES AND PARTS

We looked at online availability of parts and products of jaguar (*Panthera onca*). We were cognizant that jaguar material and postings may coincide with those of puma (*Puma concolor*), ocelot (*Leopardus pardalis*), margay (*Leopardus wiedii*), and oncilla (*Leopardus tigrinus*) in the New World. Preliminary analyses suggested that trade in the smaller cats tends towards live specimens (pets). There were indications that markets in puma and jaguar teeth overlap, as we identified through posts appearing to show both species. However, the primary focus of the grant and project was jaguar. We expected to encounter records of trade in lion (*Panthera leo*) and tiger (*Panthera tigris* spp.) in some geography/language combinations, and recorded all in standardized data sheets.

We used dynamic search terms translated into a variety of languages; starting with a list of over 200 terms, eventually refining which combinations of search terms revealed trade posts.

### MEASUREMENT OF EFFORT

We standardized the units of effort through:
1) Recording each unique search, per person, with date, person, time start and finished per search session, search term used, platform searched, productive returns, unproductive returns. Doing this can refine surveys through identifying those terms with the highest productivity.
2) An individual checklist per researcher of all combinations of search terms/body parts/platforms during each standardized search interval.
Following the above combination of practices can refine which combinations of search terms and platforms are the most effective at locating trade. However, an unintended consequence of publishing exact search terms could be that online traders deploy other terms to circumvent detection, therefore to mitigate this risk we will not publish search terms.

### POSTINGS

We defined an online post or posting as one unique post on a platform; an example is a posting containing an image of jaguar teeth. If the platform allowed for threads or comments, an associated comment related to the original posting showing an additional image was counted as a separate posting. In other words, each record is an event, much as in camera trapping (photo taken at X location and time), with contents being specified (e.g., in a metaphorical sense, one jaguar, or three peccaries). Each event was one row in an Excel, with variation according to what was encountered, e.g., one skull versus 23 teeth.

If re-listed items were found, then we recorded them as a new record and flagged them as a re-listed (duplicate) item with reference to the earlier listing/s through the unique record ID applied during data collection, for later exclusion.

**SEARCH CRITERIA AND SAMPLING**

At first, we systematically searched platforms in the suitable domains for each country/region/language.

For search results number 90-100 we reviewed if two more records were relevant within that sub-set. In that case, we continued sampling for another 10 results at a time until one in ten or fewer (e.g., zero) were relevant.

Considering all results and whether results varied across platforms, we decided whether to continue using all platforms or fewer. This happened after a first comprehensive review, not ad hoc individual decisions.

Upon returning all 100 initial search engine results, we screened indexed records and the platform names to ascertain relevance (for example, 'Jaguar' cars referred to car sales websites). If a description of the platform was unavailable, or if the nature of the platform was unclear or suggestive of positive trade results, we visited all links to ascertain what is relevant and on which platforms relevant postings appeared.

We had a set of researchers searching in Spanish, from Bolivia, and Peru (comprising the South America region) and Guatemala (comprising the Mesoamerica and Mexico region). Whilst initial searches were not restricted to these regions, when in-Spanish observers in one region encountered a record relevant to other region (such as an e-commerce site based in the other region), they recorded the basics of the productive link and shared to the relevant other regional team to complete to streamline.

**RECORDING FINDINGS**

We used a standard data template to record findings limited to project personnel only. Any changes to the data model necessitated by needs identified during the project were disseminated to the group to update their own copy.

In addition, we required that each search is accompanied by a data sheet that recorded terms, platform, number of productive/unproductive returns.

All individual searches submitted findings in two-four-week intervals for review and compilation in a centralised sheet. We conducted a detailed review of progress at the mid-way mark and refined for remainder of the period and the routine going forward after.

We counted the number of posts and the number of cat parts per post. In cases where only part of a consignment was shown, we counted what was visible and/or expressed in accompanying text as a minimum count. Text-only posts were recorded but parts not counted.

We captured suspected fakes and segmented these within our database, along with an assessment of what was fake. The expert panel reviewed this initial assessment and fakes were excluded.

We referenced records with standardized indexing.

**POTENTIAL DATA BIAS**

We attempted to avoid data biases in searching introduced by cookies stored on devices by removing cookies and browsing histories prior to each search session.

We noted that online platforms may replicate results based on previous searches, a potential bias in the results which we minimised through using as an initial search for 100 records as opposed to continuous searching.

We recognised that different jurisdictions have different levels of access to the internet which can result in variance in online content, and we used relevant national level search engines or site country domain identifiers in search terms to obtain results for appropriate jurisdictions.

We attempted to capture when postings described parts as old, heirloom or captive-source. We were aware that presence of teeth in trade is not necessarily indicative of current poaching, as an unquantified number of old teeth including those inherited and then sold may enter the trade, as may parts of captive cats.
Our 10-year search criteria may have had bias towards platforms which archive content and therefore revealed older content.

We attempted to standardise our approach as much as possible to minimise variability, noting that some variance in search terms were necessary due to language differences.

We captured when payment methods were stated, but in many cases, this may not be available information.

**EXCLUSIONS**

We did not look at the Dark Web.

In some cases, images were deemed inauthentic in themselves, although the posting might have still been a genuine offer of sale, therefore we captured our assessment in free-text within our data model, as follows:

- Stock images (widely used or potentially identifiable through a ‘reverse image search’)
- Blurry / out of focus
- Image did not match description
- Image looked old.

**PRIVACY, ETHICS AND INFORMATION GOVERNANCE**

We were cognisant of relevant national legislation related to data protection (including provisions related to processing and transfers of data) and cybercrime in addition to human rights and right to privacy. We ensured our activities and any information collection to be:

1. For non-commercial purposes
2. In proportion to the project purpose, to support efforts to better detect online jaguar and other wildlife trade
3. To capture only information relevant to the project purpose and not use tools which might collaterally identify excessive or irrelevant information
4. If posts published such information, to not capture sensitive personal data about internet users (including but not limited to information about internet users’ political affiliations or religious beliefs or health) or persons such as children or vulnerable adults
5. To identify posts only suspected of committing trade in jaguar parts
6. To maintain records which were accurate and up to date
7. To maintain records for only the time needed to fulfil the project purpose
8. To maintain records capable of being stored, reviewed and disposed of safely
9. To maintain records with access restrictions
10. To only maintain identifiers (such as URLs to trade posts) as needed for the project purpose and capable of being securely deleted or fully anonymized.

We did not make inquiries about price or product availability, which could inadvertently stimulate trade. We did not purchase or make bids on any products. Dependent on platforms, images and even the text within a posting could be considered to be under copyright, for which we considered implications of research and recording methods.

Some national jurisdictions provided a legal obligation to report crime, so where applicable we reported suspected trade posts, within the above considerations and in communications with appropriate handling and dissemination markings and confidentiality controls.
