## Supplementary material for "Multi-lingual multi-platform investigations of online trade in jaguar parts": S2 - Dataset used for results analysis

| Language of research | Platform type for search | Platform showing the post | Platform code | Year of post | Location of post (country) [ASSESSED] | If offer to transport: transport method | Telephone number displayed | Other details | Payment details displayed | Species 1 | Species 1 part/product | Species 1 number | Species 1 weight kg | Species 2 | Species 2 part/product | Species 2 number | Species 2 weight kg | Post states from captive source? | Post states old / inherited / heirloom? | Motivation for post [ASSESSED] | Image classification: A definitely jaguar, B ambiguous, C not jaguar [SPECIES 1 ONLY] | Platform type classification |
|---|---|---|---|---|---|---|---|---|---|---|---|---|---|---|---|---|---|---|---|---|---|---|
| Spanish | Search engine | Online-marketplace-13 | OM-13 | 2019 | Peru | [NOT SPECIFI | [NOT DISPLA | [NOT DISPLA | [NOT DISPLA | Jaguar | SKI Skins | 1 | | | | | | | YES | Trade | A | Online marketplace |
| Spanish | Search engine | Weblog-01 | WL-01 | 2012 | Peru | [NOT SPECIFI | [NOT DISPLA | [NOT DISPLA | [NOT DISPLA | Jaguar | SKS Skin scraps | 1 | | | | | | | | Trade | B | Weblog |
| Spanish | Social network | Social-network-01 | SN-01 | 2017 | Peru | [NOT SPECIFI | [NOT DISPLA | [NOT DISPLA | [NOT DISPLA | Jaguar | SKI Skins | 1 | | | | | | | | Trade | C | Social network |
| Spanish | Social network | Social-network-01 | SN-01 | 2016 | Peru | [NOT SPECIFI | [NOT DISPLA | [NOT DISPLA | [NOT DISPLA | Jaguar | SKI Skins | 1 | | | | | | | | Trade | A | Social network |
| Spanish | Social network | Social-network-01 | SN-01 | 2016 | Peru | [NOT SPECIFI | [NOT DISPLA | [NOT DISPLA | [NOT DISPLA | Jaguar | SKI Skins | 1 | | | | | | | | Trade | A | Social network |
| Spanish | Social network | Social-network-01 | SN-01 | 2017 | Peru | [NOT SPECIFI | [NOT DISPLA | [NOT DISPLA | [NOT DISPLA | Jaguar | SKI Skins | 1 | | | | | | | | Trade | A | Social network |
| Spanish | Social network | Social-network-01 | SN-01 | 2015 | Peru | [NOT SPECIFI | DISPLAYED | [NOT DISPLA | [NOT DISPLA | Jaguar | SKI Skins | 1 | | | | | | | | Trade | A | Social network |
| Spanish | Social network | Social-network-01 | SN-01 | 2017 | Peru | [NOT SPECIFI | [NOT DISPLA | [NOT DISPLA | [NOT DISPLA | Jaguar | SKI Skins | 1 | | | | | | | | Trade | A | Social network |
| French | Search engine | Online-marketplace-08 | OM-08 | [NOT SPECIFI | Guadeloupe | [NOT SPECIFI | [NOT DISPLA | [NOT DISPLA | [NOT DISPLA | Jaguar | TEE Teeth | 1 | | | | | | | | Trade | A | Online marketplace |
| Vietnamese | Search engine | Video-sharing-02 | VS-02 | 2019 | Viet Nam | [NOT SPECIFI | DISPLAYED | ADDITIONAL | CONTACT SE | Jaguar | TEE Teeth | 9 | | Tiger | CLA Claws | 2 | | | | Trade | C | Video-sharing |
| Vietnamese | Social network | Social-network-05 | SN-05 | 2019 | Viet Nam | BY POST | DISPLAYED | ADDRESS DIS | TRANSFER SE | Jaguar | TEE Teeth | 1 | | Tiger | TEE Teeth | 4 | | | | Trade | C | Social network |
| Vietnamese | Search engine | Video-sharing-02 | VS-02 | 2018 | Viet Nam | [NOT SPECIFI | DISPLAYED | ADDITIONAL | [NOT DISPLA | Jaguar | TEE Teeth | 1 | | | | | | | | Trade | C | Video-sharing |
| Vietnamese | Search engine | Video-sharing-02 | VS-02 | 2019 | Viet Nam | [NOT SPECIFI | DISPLAYED | ADDITIONAL | [NOT DISPLA | Jaguar | TEE Teeth | 4 | | Tiger | CLA Claws | 2 | | YES | | Trade | C | Video-sharing |
| Vietnamese | Search engine | Social-network-05 | SN-05 | 2018 | Viet Nam | [NOT SPECIFI | DISPLAYED | ADDITIONAL | [NOT DISPLA | Jaguar | TEE Teeth | 6 | | Tiger | CLA Claws | 1 | | | | Trade | C | Social network |
| Vietnamese | Search engine | Video-sharing-01 | VS-01 | 2018 | Viet Nam | [NOT SPECIFI | DISPLAYED | ADDITIONAL | [NOT DISPLA | Jaguar | TEE Teeth | 2 | | Tiger | TEE Teeth | 2 | | | | Trade | C | Video-sharing |
| Vietnamese | Search engine | Video-sharing-01 | VS-01 | 2019 | Viet Nam | [NOT SPECIFI | DISPLAYED | ADDITIONAL | [NOT DISPLA | Jaguar | TEE Teeth | 2 | | | | | | | | Trade | C | Video-sharing |
| Vietnamese | Search engine | Video-sharing-01 | VS-01 | 2019 | Viet Nam | [NOT SPECIFI | DISPLAYED | ADDITIONAL | [NOT DISPLA | Jaguar | TEE Teeth | 1 | | | | | | | | Trade | C | Video-sharing |
| Vietnamese | Search engine | Video-sharing-01 | VS-01 | 2018 | Viet Nam | [NOT SPECIFI | DISPLAYED | ADDITIONAL | [NOT DISPLA | Jaguar | TEE Teeth | 5 | | | | | | | | Trade | C | Video-sharing |
| Vietnamese | Search engine | Video-sharing-01 | VS-01 | 2018 | Viet Nam | [NOT SPECIFI | DISPLAYED | [NOT DISPLA | [NOT DISPLA | Jaguar | TEE Teeth | 1 | | Tiger | TEE Teeth | 1 | | | | Trade | C | Video-sharing |
| Vietnamese | Search engine | Video-sharing-01 | VS-01 | 2018 | Viet Nam | [NOT SPECIFI | DISPLAYED | ADDITIONAL | [NOT DISPLA | Jaguar | TEE Teeth | 2 | | Tiger | CLA Claws | 8 | | | | Trade | C | Video-sharing |
| Vietnamese | Search engine | Video-sharing-01 | VS-01 | 2018 | Viet Nam | [NOT SPECIFI | DISPLAYED | ADDITIONAL | [NOT DISPLA | Jaguar | TEE Teeth | 2 | | | | | | | | Trade | C | Video-sharing |
| Vietnamese | Search engine | Video-sharing-01 | VS-01 | 2019 | Viet Nam | [NOT SPECIFI | DISPLAYED | ADDITIONAL | [NOT DISPLA | Jaguar | TEE Teeth | 8 | | Tiger | CLA Claws | 10 | | | | Trade | B | Video-sharing |
| Vietnamese | Search engine | Video-sharing-01 | VS-01 | 2018 | Viet Nam | [NOT SPECIFI | DISPLAYED | ADDITIONAL | [NOT DISPLA | Jaguar | TEE Teeth | 2 | | | | | | | | Trade | C | Video-sharing |
| Vietnamese | Search engine | Video-sharing-01 | VS-01 | 2018 | Viet Nam | [NOT SPECIFI | DISPLAYED | ADDITIONAL | [NOT DISPLA | Jaguar | TEE Teeth | 1 | | | | | | | | Trade | UNCLEAR-POSSIBLY B | Video-sharing |
| Vietnamese | Search engine | Video-sharing-01 | VS-01 | 2018 | Viet Nam | [NOT SPECIFI | DISPLAYED | ADDITIONAL | [NOT DISPLA | Jaguar | TEE Teeth | 1 | | | | | | | | Trade | C | Video-sharing |
| Vietnamese | Search engine | Video-sharing-01 | VS-01 | 2019 | Viet Nam | [NOT SPECIFI | DISPLAYED | ADDITIONAL | [NOT DISPLA | Jaguar | TEE Teeth | 3 | | Tiger | TEE Teeth | 2 | | | | Trade | C | Video-sharing |
| Vietnamese | Search engine | Video-sharing-01 | VS-01 | 2019 | Viet Nam | [NOT SPECIFI | DISPLAYED | [NOT DISPLA | [NOT DISPLA | Jaguar | TEE Teeth | 9 | | Tiger | CLA Claws | 2 | | | | Trade | C | Video-sharing |
| Vietnamese | Search engine | Video-sharing-01 | VS-01 | 2019 | Viet Nam | [NOT SPECIFI | DISPLAYED | ADDITIONAL | [NOT DISPLA | Jaguar | TEE Teeth | 3 | | | | | | | | Trade | C | Video-sharing |
| Vietnamese | Search engine | Video-sharing-01 | VS-01 | 2018 | Viet Nam | [NOT SPECIFI | DISPLAYED | ADDITIONAL | [NOT DISPLA | Jaguar | TEE Teeth | 2 | | | | | | | | Trade | C | Video-sharing |
| Vietnamese | Search engine | Video-sharing-01 | VS-01 | 2019 | Viet Nam | [NOT SPECIFI | DISPLAYED | ADDITIONAL | [NOT DISPLA | Jaguar | TEE Teeth | 6 | | Tiger | CLA Claws | 3 | | | | Trade | B | Video-sharing |
| Vietnamese | Search engine | Video-sharing-01 | VS-01 | 2018 | Viet Nam | [NOT SPECIFI | DISPLAYED | ADDITIONAL | [NOT DISPLA | Jaguar | TEE Teeth | 3 | | | | | | | | Trade | C | Video-sharing |
| Vietnamese | Search engine | Video-sharing-01 | VS-01 | 2018 | Viet Nam | [NOT SPECIFI | DISPLAYED | ADDITIONAL | [NOT DISPLA | Jaguar | TEE Teeth | 4 | | | | | | | | Trade | C | Video-sharing |
| Vietnamese | Search engine | Video-sharing-01 | VS-01 | 2018 | Viet Nam | [NOT SPECIFI | DISPLAYED | ADDITIONAL | [NOT DISPLA | Jaguar | TEE Teeth | 1 | | | | | | | | Trade | C | Video-sharing |
| Vietnamese | Search engine | Video-sharing-01 | VS-01 | 2018 | Viet Nam | [NOT SPECIFI | DISPLAYED | ADDITIONAL | [NOT DISPLA | Jaguar | TEE Teeth | 3 | | Tiger | CLA Claws | 1 | | | | Trade | C | Video-sharing |
| Vietnamese | Search engine | Video-sharing-01 | VS-01 | 2018 | Viet Nam | [NOT SPECIFI | DISPLAYED | ADDITIONAL | [NOT DISPLA | Jaguar | TEE Teeth | 4 | | | | | | | | Trade | B | Video-sharing |
| Vietnamese | Search engine | Video-sharing-01 | VS-01 | 2018 | Viet Nam | [NOT SPECIFI | DISPLAYED | ADDITIONAL | [NOT DISPLA | Jaguar | TEE Teeth | 1 | | | | | | | | Trade | C | Video-sharing |
| Vietnamese | Social network | Social-network-01 | SN-01 | 2014 | Viet Nam | [NOT SPECIFI | [NOT DISPLA | [NOT DISPLA | [NOT DISPLA | Jaguar | TEE Teeth | 1 | | | | | | | | Trade | C | Social network |
| Vietnamese | Social network | Social-network-01 | SN-01 | 2019 | Viet Nam | [NOT SPECIFI | [NOT DISPLA | [NOT DISPLA | [NOT DISPLA | Jaguar | TEE Teeth | 1 | | | | | | | | Trade | B | Social network |
| Vietnamese | Social network | Social-network-01 | SN-01 | 2019 | Viet Nam | [NOT SPECIFI | [NOT DISPLA | [NOT DISPLA | [NOT DISPLA | Jaguar | TEE Teeth | 10 | | Tiger | TEE Teeth | 4 | | | | Trade | C | Social network |
| Vietnamese | Social network | Social-network-01 | SN-01 | 2019 | Viet Nam | [NOT SPECIFI | [NOT DISPLA | [NOT DISPLA | [NOT DISPLA | Jaguar | TEE Teeth | 1 | | Tiger | TEE Teeth | 4 | | | | Trade | C | Social network |
| Spanish | Social network | Social-network-01 | SN-01 | 2014 | Mexico | [NOT SPECIFI | [NOT DISPLA | [NOT DISPLA | [NOT DISPLA | Jaguar | TEE Teeth | 1 | | Puma | TEE Teeth | 2 | | | | Trade | A | Social network |
| Spanish | Social network | Social-network-01 | SN-01 | 2015 | Mexico | [NOT SPECIFI | [NOT DISPLA | [NOT DISPLA | [NOT DISPLA | Jaguar | TEE Teeth | 2 | | | | | | | | Trade | A | Social network |
| Spanish | Social network | Social-network-01 | SN-01 | 2017 | Mexico | [NOT SPECIFI | [NOT DISPLA | [NOT DISPLA | [NOT DISPLA | Jaguar | TEE Teeth | 1 | | | | | | | | Trade | B | Social network |
| Spanish | Social network | Social-network-01 | SN-01 | 2015 | Mexico | [NOT SPECIFI | [NOT DISPLA | [NOT DISPLA | [NOT DISPLA | Jaguar | TEE Teeth | 1 | | | | | | | | Trade | A | Social network |
| Spanish | Social network | Social-network-01 | SN-01 | 2018 | Mexico | [NOT SPECIFI | [NOT DISPLA | [NOT DISPLA | CONTACT SE | Jaguar | TEE Teeth | 3 | | | | | | | | Trade | B | Social network |
| Spanish | Social network | Social-network-01 | SN-01 | 2017 | Mexico | [NOT SPECIFI | DISPLAYED | [NOT DISPLA | [NOT DISPLA | Jaguar | TEE Teeth | 1 | | | | | | | | Trade | A | Social network |
| Spanish | Social network | Social-network-01 | SN-01 | 2015 | Mexico | [NOT SPECIFI | [NOT DISPLA | [NOT DISPLA | [NOT DISPLA | Jaguar | TEE Teeth | 3 | | | | | | | | Trade | A | Social network |
| Spanish | Social network | Social-network-01 | SN-01 | 2019 | Mexico | [NOT SPECIFI | [NOT DISPLA | [NOT DISPLA | [NOT DISPLA | Jaguar | TEE Teeth | 1 | | | | | | | | Trade | B | Social network |
| Spanish | Social network | Social-network-01 | SN-01 | 2014 | Mexico | [NOT SPECIFI | [NOT DISPLA | [NOT DISPLA | [NOT DISPLA | Jaguar | TEE Teeth | 1 | | | | | | | | Trade | A | Social network |
| Spanish | Social network | Social-network-01 | SN-01 | 2016 | Mexico | [NOT SPECIFI | [NOT DISPLA | [NOT DISPLA | [NOT DISPLA | Jaguar | TEE Teeth | 3 | | | | | | | | Trade | A | Social network |
| Spanish | Social network | Social-network-01 | SN-01 | 2013 | Mexico | [NOT SPECIFI | [NOT DISPLA | [NOT DISPLA | [NOT DISPLA | Jaguar | TEE Teeth | 2 | | | | | | | | Trade | B | Social network |
| Spanish | Social network | Social-network-01 | SN-01 | 2014 | Mexico | [NOT SPECIFI | [NOT DISPLA | [NOT DISPLA | [NOT DISPLA | Jaguar | TEE Teeth | 1 | | Puma | TEE Teeth | 4 | | | | Trade | A | Social network |
| Spanish | Social network | Social-network-01 | SN-01 | 2019 | Mexico | [NOT SPECIFI | [NOT DISPLA | [NOT DISPLA | [NOT DISPLA | Jaguar | TEE Teeth | 1 | | | | | | | | Trade | A | Social network |
| Spanish | Search engine | Online-marketplace-13 | OM-13 | 2018 | Mexico | [NOT SPECIFI | [NOT DISPLA | [NOT DISPLA | [NOT DISPLA | Jaguar | CLA Claws | 1 | | | | | | | | Trade | B | Online marketplace |
| Spanish | Search engine | Online-marketplace-13 | OM-13 | [NOT SPECIFI | Mexico | [NOT SPECIFI | [NOT DISPLA | [NOT DISPLA | [NOT DISPLA | Jaguar | CLA Claws | 1 | | | | | | | | Trade | B | Online marketplace |
| Spanish | Search engine | Online-marketplace-13 | OM-13 | 2019 | Mexico | [NOT SPECIFI | [NOT DISPLA | [NOT DISPLA | [NOT DISPLA | Jaguar | CLA Claws | 1 | | | | | | | | Trade | B | Online marketplace |
| Spanish | Search engine | Search-engine-02 | SE-02 | 2016 | Mexico | [NOT SPECIFI | DISPLAYED | [NOT DISPLA | [NOT DISPLA | Jaguar | SKI Skins | 1 | | | | | | | | Trade | UNCLEAR-POSSIBLY C | Search engine |
| Spanish | Social network | Social-network-01 | SN-01 | 2013 | Mexico | [NOT SPECIFI | [NOT DISPLA | [NOT DISPLA | [NOT DISPLA | Jaguar | SKI Skins | 1 | | | | | | | | Trade | C | Social network |
| Spanish | Social network | Social-network-01 | SN-01 | 2013 | Mexico | [NOT SPECIFI | [NOT DISPLA | [NOT DISPLA | [NOT DISPLA | Jaguar | SKI Skins | 1 | | | | | | | | Trade | A | Social network |
| Spanish | Social network | Social-network-01 | SN-01 | 2012 | Mexico | [NOT SPECIFI | [NOT DISPLA | [NOT DISPLA | [NOT DISPLA | Jaguar | TEE Teeth | 1 | | | | | | | | Trade | B | Social network |
| Spanish | Social network | Social-network-01 | SN-01 | 2013 | Mexico | [NOT SPECIFI | [NOT DISPLA | [NOT DISPLA | [NOT DISPLA | Jaguar | TEE Teeth | 1 | | | | | | | | Trade | B | Social network |
| Spanish | Social network | Social-network-01 | SN-01 | 2013 | Uruguay | [NOT SPECIFI | DISPLAYED | [NOT DISPLA | [NOT DISPLA | Jaguar | TEE Teeth | 1 | | | | | | | | Trade | A | Social network |
| Spanish | Social network | Social-network-01 | SN-01 | 2015 | Mexico | [NOT SPECIFI | [NOT DISPLA | [NOT DISPLA | [NOT DISPLA | Jaguar | CLA Claws | 2 | | Shark | TEE Teeth | 1 | | | | Trade | B | Social network |
| Spanish | Social network | Social-network-01 | SN-01 | 2017 | Costa Rica | [NOT SPECIFI | [NOT DISPLA | [NOT DISPLA | [NOT DISPLA | Jaguar | CLA Claws | 1 | | Sea Lion | TEE Teeth | 1 | | | | Trade | B | Social network |
| Spanish | Social network | Social-network-01 | SN-01 | 2015 | Mexico | [NOT SPECIFI | [NOT DISPLA | [NOT DISPLA | [NOT DISPLA | Jaguar | TEE Teeth | 1 | | | | | | | | Trade | B | Social network |
| Spanish | Social network | Social-network-01 | SN-01 | 2017 | Mexico | [NOT SPECIFI | [NOT DISPLA | [NOT DISPLA | [NOT DISPLA | Jaguar | TEE Teeth | 1 | | | | | | | | Trade | C | Social network |
| Spanish | Social network | Social-network-01 | SN-01 | 2017 | Mexico | [NOT SPECIFI | [NOT DISPLA | [NOT DISPLA | [NOT DISPLA | Jaguar | TEE Teeth | 1 | | | | | | | | Trade | C | Social network |
| Spanish | Social network | Social-network-01 | SN-01 | 2015 | Mexico | [NOT SPECIFI | [NOT DISPLA | [NOT DISPLA | [NOT DISPLA | Jaguar | TEE Teeth | 1 | | | | | | | | Trade | B | Social network |
| Spanish | Social network | Social-network-01 | SN-01 | 2018 | Mexico | BY COURIER | [NOT DISPLA | [NOT DISPLA | [NOT DISPLA | Jaguar | TEE Teeth | 1 | | | | | | | | Trade | A | Social network |
| Spanish | Social network | Social-network-01 | SN-01 | 2017 | Mexico | [NOT SPECIFI | [NOT DISPLA | [NOT DISPLA | [NOT DISPLA | Jaguar | TEE Teeth | 4 | | | | | | | | Trade | A | Social network |
| Spanish | Social network | Social-network-01 | SN-01 | 2017 | Mexico | [NOT SPECIFI | [NOT DISPLA | [NOT DISPLA | [NOT DISPLA | Jaguar | TEE Teeth | 10 | | Tiger | TEE Teeth | 4 | | | | Trade | B | Social network |
| Spanish | Social network | Social-network-01 | SN-01 | 2017 | Mexico | [NOT SPECIFI | [NOT DISPLA | [NOT DISPLA | [NOT DISPLA | Jaguar | TEE Teeth | 1 | | | | | | | | Trade | B | Social network |
| Spanish | Social network | Social-network-01 | SN-01 | 2018 | Mexico | [NOT SPECIFI | [NOT DISPLA | [NOT DISPLA | [NOT DISPLA | Jaguar | TEE Teeth | 1 | | | | | | | | Trade | B | Social network |
| Spanish | Social network | Social-network-01 | SN-01 | 2019 | Mexico | [NOT SPECIFI | [NOT DISPLA | [NOT DISPLA | [NOT DISPLA | Jaguar | TEE Teeth | 1 | | | | | | | | Trade | B | Social network |
| Spanish | Social network | Social-network-01 | SN-01 | 2019 | Guatemala | [NOT SPECIFI | [NOT DISPLA | [NOT DISPLA | [NOT DISPLA | Jaguar | TEE Teeth | 1 | | | | | | | | Trade | B | Social network |
| Spanish | Social network | Social-network-01 | SN-01 | 2014 | Mexico | [NOT SPECIFI | [NOT DISPLA | [NOT DISPLA | [NOT DISPLA | Jaguar | TEE Teeth | 1 | | | | | | | | Trade | B | Social network |
| Spanish | Search engine | Search-engine-02 | SE-02 | 2010 | Mexico | [NOT SPECIFI | [NOT DISPLA | [NOT DISPLA | [NOT DISPLA | Jaguar | TEE Teeth | 1 | | | | | | | YES | Trade | B | Search engine |
| Spanish | Search engine | Online-marketplace-13 | OM-13 | 2017 | Mexico | [NOT SPECIFI | [NOT DISPLA | [NOT DISPLA | [NOT DISPLA | Jaguar | TEE Teeth | 1 | | | | | | | | Trade | [NO IMAGE REVIEW] | Online marketplace |
| Spanish | Search engine | Online-marketplace-07 | OM-07 | 2016 | Mexico | [NOT SPECIFI | [NOT DISPLA | [NOT DISPLA | [NOT DISPLA | Jaguar | TEE Teeth | 1 | | | | | | | YES | Trade | B | Online marketplace |
| Spanish | Search engine | Online-marketplace-13 | OM-13 | 2018 | Mexico | [NOT SPECIFI | [NOT DISPLA | [NOT DISPLA | [NOT DISPLA | Jaguar | CLA Claws | 1 | | | | | | | | Trade | [NO IMAGE REVIEW] | Online marketplace |
| Spanish | Social network | Social-network-01 | SN-01 | 2014 | Mexico | [NOT SPECIFI | [NOT DISPLA | [NOT DISPLA | [NOT DISPLA | Jaguar | CLA Claws | 2 | | | | | | | | Trade | B | Social network |
| Spanish | Social network | Social-network-01 | SN-01 | 2017 | Mexico | [NOT SPECIFI | [NOT DISPLA | [NOT DISPLA | [NOT DISPLA | Jaguar | CLA Claws | 1 | | | | | | | | Trade | B | Social network |
| Spanish | Social network | Social-network-01 | SN-01 | 2015 | Mexico | [NOT SPECIFI | [NOT DISPLA | [NOT DISPLA | [NOT DISPLA | Jaguar | CLA Claws | 2 | | Peccary | TEE Teeth | 2 | | | | Trade | B | Social network |
| Spanish | Social network | Social-network-01 | SN-01 | 2015 | Mexico | [NOT SPECIFI | [NOT DISPLA | [NOT DISPLA | [NOT DISPLA | Jaguar | SKI Skins | 1 | | | | | | | | Trade | A | Social network |
| Spanish | Social network | Social-network-01 | SN-01 | 2017 | Mexico | [NOT SPECIFI | [NOT DISPLA | [NOT DISPLA | [NOT DISPLA | Jaguar | SKU Skulls | 1 | | | | | | | | Trade | B | Social network |
| Spanish | Social network | Social-network-01 | SN-01 | 2017 | Mexico | [NOT SPECIFI | [NOT DISPLA | [NOT DISPLA | [NOT DISPLA | Jaguar | CLA Claws | 1 | | | | | | | | Trade | B | Social network |
| Spanish | Social network | Social-network-01 | SN-01 | 2015 | Mexico | [NOT SPECIFI | [NOT DISPLA | [NOT DISPLA | [NOT DISPLA | Jaguar | BOP Bone pieces | 1 | | | | | | | | Trade | C | Social network |
| Spanish | Social network | Social-network-01 | SN-01 | 2013 | Mexico | [NOT SPECIFI | [NOT DISPLA | [NOT DISPLA | [NOT DISPLA | Jaguar | SKP Skin pieces | 1 | | Crocodile | SKP Skin piec | 1 | | | | Trade | A | Social network |
| Spanish | Social network | Social-network-01 | SN-01 | 2018 | Nicaragua | [NOT SPECIFI | [NOT DISPLA | [NOT DISPLA | [NOT DISPLA | Jaguar | SKI Skins | 1 | | | | | | | | Trade | A | Social network |
| Spanish | Search engine | Social-network-04 | SN-04 | 2015 | Colombia | [NOT SPECIFI | [NOT DISPLA | [NOT DISPLA | [NOT DISPLA | Jaguar | SKI Skins | 1 | | | | | | | INHERITED | Trade | [NO IMAGE REVIEW] | Social network |
| Spanish | Search engine | Online-marketplace-13 | OM-13 | 2019 | Ecuador | [NOT SPECIFI | [NOT DISPLA | [NOT DISPLA | CONTACT SE | Jaguar | TEE Teeth | 1 | | | | | | | | Trade | C | Online marketplace |
| Spanish | Search engine | Online-marketplace-13 | OM-13 | 2019 | Venezuela | BUYER COLLI | [NOT DISPLA | [NOT DISPLA | [NOT DISPLA | Jaguar | HEA Heads | 1 | | | | | | | YES | Trade | A | Online marketplace |
| Spanish | Search engine | Online-marketplace-13 | OM-13 | 2018 | Argentina | [PUBLISHED | [NOT DISPLA | [NOT DISPLA | CASH, CREDI | Jaguar | TEE Teeth | 1 | | | | | | | | Trade | B | Online marketplace |
| Spanish | Search engine | Online-marketplace-13 | OM-13 | 2019 | Venezuela | TO BE AGREI | [NOT DISPLA | [NOT DISPLA | CONTACT SE | Jaguar | SKI Skins | 1 | | | | | | | YES | Trade | A | Online marketplace |
| Spanish | Social network | Social-network-01 | SN-01 | 2019 | Venezuela | [NOT SPECIFI | [NOT DISPLA | [NOT DISPLA | [NOT DISPLA | Jaguar | TEE Teeth | 3 | | | | | | | | Trade | A | Social network |
| Spanish | Social network | Social-network-01 | SN-01 | 2017 | Venezuela | [NOT SPECIFI | DISPLAYED | [NOT DISPLA | [NOT DISPLA | Jaguar | SKI Skins | 1 | | | | | | | | Trade | C | Social network |
| Portuguese | Social network | Social-network-01 | SN-01 | 2017 | Brazil | [NOT SPECIFI | DISPLAYED | [NOT DISPLA | [NOT DISPLA | Jaguar | TEE Teeth | 2 | | | | | | | | Trade | B | Social network |
| Portuguese | Social network | Social-network-01 | SN-01 | 2017 | Brazil | [NOT SPECIFI | DISPLAYED | [NOT DISPLA | [NOT DISPLA | Jaguar | TEE Teeth | 1 | | | | | | | | Trade | A | Social network |
| Portuguese | Social network | Social-network-01 | SN-01 | 2018 | Brazil | [NOT SPECIFI | DISPLAYED | [NOT DISPLA | [NOT DISPLA | Jaguar | TEE Teeth | 1 | | | | | | | | Trade | B | Social network |
| Portuguese | Social network | Social-network-01 | SN-01 | 2015 | Brazil | [NOT SPECIFI | DISPLAYED | [NOT DISPLA | [NOT DISPLA | Jaguar | TEE Teeth | 1 | | | | | | | | Trade | C | Social network |
| Portuguese | Social network | Social-network-01 | SN-01 | 2018 | Brazil | BY POST | [NOT DISPLA | [NOT DISPLA | [NOT DISPLA | Jaguar | TEE Teeth | 1 | | | | | | | | Trade | A | Social network |
| Portuguese | Social network | Social-network-01 | SN-01 | 2019 | Brazil | [NOT SPECIFI | [NOT DISPLA | [NOT DISPLA | [NOT DISPLA | Jaguar | TEE Teeth | 1 | | | | | | | | Trade | [NO IMAGE REVIEW] | Social network |
| Portuguese | Social network | Social-network-01 | SN-01 | 2015 | Brazil | BY POST | [NOT DISPLA | [NOT DISPLA | [NOT DISPLA | Jaguar | TEE Teeth | 1 | | | | | | | | Trade | A | Social network |
| Portuguese | Social network | Social-network-01 | SN-01 | 2015 | Brazil | [NOT SPECIFI | [NOT DISPLA | [NOT DISPLA | [NOT DISPLA | Jaguar | TEE Teeth | 1 | | | | | | | | Trade | C | Social network |
| Portuguese | Social network | Social-network-01 | SN-01 | 2015 | Brazil | [NOT SPECIFI | DISPLAYED | [NOT DISPLA | [NOT DISPLA | Jaguar | TEE Teeth | 3 | | | | | | | | Trade | C | Social network |
| Portuguese | Social network | Social-network-01 | SN-01 | 2016 | Brazil | [NOT SPECIFI | DISPLAYED | [NOT DISPLA | [NOT DISPLA | Jaguar | TEE Teeth | 1 | | Puma | CLA Claws | 1 | | | | Trade | C | Social network |
| Portuguese | Social network | Social-network-01 | SN-01 | 2019 | Brazil | [NOT SPECIFI | [NOT DISPLA | [NOT DISPLA | [NOT DISPLA | Jaguar | TEE Teeth | 1 | | | | | | | | Trade | B | Social network |
| Portuguese | Social network | Social-network-01 | SN-01 | 2017 | Brazil | [NOT SPECIFI | DISPLAYED | [NOT DISPLA | [NOT DISPLA | Jaguar | TEE Teeth | 1 | | | | | | | | Trade | A | Social network |
| Portuguese | Social network | Social-network-01 | SN-01 | 2018 | Brazil | [NOT SPECIFI | [NOT DISPLA | [NOT DISPLA | [NOT DISPLA | Jaguar | TEE Teeth | 1 | | | | | | | | Trade | A | Social network |
| Portuguese | Social network | Social-network-01 | SN-01 | 2017 | Brazil | [NOT SPECIFI | [NOT DISPLA | [NOT DISPLA | [NOT DISPLA | Jaguar | TEE Teeth | 1 | | | | | | | | Trade | A | Social network |
| Portuguese | Social network | Social-network-01 | SN-01 | 2017 | Brazil | [NOT SPECIFI | [NOT DISPLA | [NOT DISPLA | [NOT DISPLA | Jaguar | TEE Teeth | 1 | | | | | | | | Trade | C | Social network |
| Portuguese | Online marketplace | Online-marketplace-03 | OM-03 | 2019 | Brazil | BY POST | [NOT DISPLA | [NOT DISPLA | [NOT DISPLA | Jaguar | SKI Skins | 1 | | | | | | | | Trade | A | Online marketplace |
| Portuguese | Online marketplace | Online-marketplace-09 | OM-09 | 2015 | Brazil | [NOT SPECIFI | DISPLAYED | [NOT DISPLA | [NOT DISPLA | Jaguar | SKI Skins | 1 | | | | | | | | Trade | B | Online marketplace |
| Portuguese | Online marketplace | Online-marketplace-06 | OM-06 | 2016 | Brazil | [NOT SPECIFI | DISPLAYED | [NOT DISPLA | [NOT DISPLA | Jaguar | SKI Skins | 1 | | | | | | | | Trade | C | Online marketplace |
| Portuguese | Online marketplace | Online-marketplace-10 | OM-10 | 2013 | Brazil | [NOT SPECIFI | DISPLAYED | [NOT DISPLA | [NOT DISPLA | Jaguar | SKI Skins | 1 | | | | | | | | Trade | A | Online marketplace |
| Portuguese | Online marketplace | Online-marketplace-10 | OM-10 | 2014 | Brazil | [NOT SPECIFI | DISPLAYED | [NOT DISPLA | [NOT DISPLA | Jaguar | SKI Skins | 1 | | | | | | | | Trade | A | Online marketplace |
| Portuguese | Online marketplace | Online-marketplace-14 | OM-14 | 2018 | Brazil | [NOT SPECIFI | DISPLAYED | [NOT DISPLA | [NOT DISPLA | Jaguar | SKI Skins | 1 | | | | | | | | Trade | A | Online marketplace |
| Portuguese | Online marketplace | Online-marketplace-10 | OM-10 | 2016 | Brazil | [NOT SPECIFI | DISPLAYED | [NOT DISPLA | [NOT DISPLA | Jaguar | SKI Skins | 1 | | | | | | | | Trade | A | Online marketplace |
| Portuguese | Online marketplace | Online-marketplace-15 | OM-15 | [NOT SPECIFI | Brazil | [NOT SPECIFI | DISPLAYED | [NOT DISPLA | [NOT DISPLA | Jaguar | SKI Skins | 1 | | | | | | | | Trade | A | Online marketplace |
| Portuguese | Online marketplace | Online-marketplace-02 | OM-02 | [NOT SPECIFI | USA | [NOT SPECIFI | DISPLAYED | [NOT DISPLA | [NOT DISPLA | Jaguar | TEE Teeth | 1 | | | | | | | | Trade | C | Online marketplace |
| Portuguese | Online marketplace | Online-marketplace-02 | OM-02 | [NOT SPECIFI | USA | [NOT SPECIFI | DISPLAYED | [NOT DISPLA | [NOT DISPLA | Jaguar | TEE Teeth | 1 | | | | | | | | Trade | C | Online marketplace |
| Portuguese | Online marketplace | Online-marketplace-05 | OM-05 | 2015 | Brazil | [NOT SPECIFI | DISPLAYED | [NOT DISPLA | [NOT DISPLA | Jaguar | TEE Teeth | 2 | | | | | | | | Trade | C | Online marketplace |
| Portuguese | Online marketplace | Online-marketplace-05 | OM-05 | 2015 | Brazil | [NOT SPECIFI | DISPLAYED | [NOT DISPLA | [NOT DISPLA | Jaguar | TEE Teeth | 2 | | | | | | | | Trade | C | Online marketplace |
| Portuguese | Social network | Social-network-01 | SN-01 | 2016 | Brazil | [NOT SPECIFI | [NOT DISPLA | [NOT DISPLA | [NOT DISPLA | Jaguar | TEE Teeth | 1 | | | | | | | | Trade | B | Social network |
| Portuguese | Social network | Social-network-01 | SN-01 | 2019 | Brazil | [NOT SPECIFI | [NOT DISPLA | [NOT DISPLA | [NOT DISPLA | Jaguar | TEE Teeth | 1 | | | | | | | | Trade | B | Social network |
| Portuguese | Social network | Social-network-01 | SN-01 | 2019 | Brazil | [NOT SPECIFI | [NOT DISPLA | [NOT DISPLA | [NOT DISPLA | Jaguar | TEE Teeth | 1 | | | | | | | | Trade | C | Social network |
| Portuguese | Social network | Social-network-01 | SN-01 | 2016 | Brazil | BY POST | DISPLAYED | [NOT DISPLA | [NOT DISPLA | Jaguar | TEE Teeth | 1 | | | | | | | | Trade | B | Social network |
| Portuguese | Online marketplace | Online-marketplace-11 | OM-11 | 2015 | Brazil | [NOT SPECIFI | DISPLAYED | [NOT DISPLA | [NOT DISPLA | Jaguar | TEE Teeth | 1 | | | | | | | | Trade | A | Online marketplace |
| Portuguese | Online marketplace | Online-marketplace-11 | OM-11 | 2015 | Brazil | [NOT SPECIFI | DISPLAYED | [NOT DISPLA | [NOT DISPLA | Jaguar | SKI Skins | 1 | | | | | | | | Trade | A | Online marketplace |
| Portuguese | Online marketplace | Online-marketplace-11 | OM-11 | 2015 | Brazil | [NOT SPECIFI | DISPLAYED | [NOT DISPLA | [NOT DISPLA | Jaguar | BOD Bodies | 1 | | | | | | | | Trade | A | Online marketplace |
| Portuguese | Online marketplace | Online-marketplace-13 | OM-13 | 2018 | Brazil | [NOT SPECIFI | [NOT DISPLA | [NOT DISPLA | [NOT DISPLA | Jaguar | TEE Teeth | 38 | | | | | | | | Trade | B | Online marketplace |
| Portuguese | Social network | Social-network-01 | SN-01 | 2018 | Brazil | [NOT SPECIFI | [NOT DISPLA | [NOT DISPLA | [NOT DISPLA | Jaguar | TEE Teeth | 1 | | | | | | | | Trade | B | Social network |
| Portuguese | Social network | Social-network-01 | SN-01 | 2016 | Brazil | [NOT SPECIFI | [NOT DISPLA | [NOT DISPLA | [NOT DISPLA | Jaguar | TEE Teeth | 1 | | | | | | | | Trade | A | Social network |
| Portuguese | Social network | Social-network-01 | SN-01 | 2016 | Brazil | [NOT SPECIFI | [NOT DISPLA | [NOT DISPLA | [NOT DISPLA | Jaguar | TEE Teeth | 1 | | | | | | | | Trade | B | Social network |
| Portuguese | Social network | Social-network-01 | SN-01 | 2017 | Brazil | [NOT SPECIFI | [NOT DISPLA | [NOT DISPLA | [NOT DISPLA | Jaguar | BOD Bodies | 1 | | | | | | | | Trade | A | Social network |
| Portuguese | Social network | Social-network-01 | SN-01 | 2015 | Brazil | [NOT SPECIFI | [NOT DISPLA | [NOT DISPLA | [NOT DISPLA | Jaguar | TEE Teeth | 1 | | | | | | | | Trade | A | Social network |
| Portuguese | Social network | Social-network-01 | SN-01 | 2017 | Brazil | [NOT SPECIFI | [NOT DISPLA | [NOT DISPLA | [NOT DISPLA | Jaguar | SKI Skins | 1 | | | | | | | | Trade | B | Social network |
| Portuguese | Social network | Social-network-01 | SN-01 | 2019 | Brazil | [NOT SPECIFI | [NOT DISPLA | [NOT DISPLA | [NOT DISPLA | Jaguar | TEE Teeth | 1 | | | | | | | | Trade | B | Social network |
| Portuguese | Social network | Social-network-01 | SN-01 | 2017 | Brazil | BY POST | DISPLAYED | [NOT DISPLA | [NOT DISPLA | Jaguar | TEE Teeth | 1 | | | | | | | | Trade | B | Social network |
| Portuguese | Social network | Social-network-01 | SN-01 | 2017 | Brazil | BY POST | DISPLAYED | [NOT DISPLA | [NOT DISPLA | Jaguar | TEE Teeth | 1 | | | | | | | | Trade | B | Social network |
| Chinese | Social network | Social-network-02 | SN-02 | 2019 | | [NOT SPECIFI | [NOT DISPLA | [NOT DISPLA | [NOT DISPLA | Jaguar | [NOT SPECIFIED] | [NOT SPECIFIED] | | | | | | | | Trade | [NO IMAGE REVIEW] | Social network |
| Chinese | Online marketplace | Online-marketplace-20 | OM-20 | 2019 | China | [NOT SPECIFI | [NOT DISPLA | [NOT DISPLA | TRANSFER SE | Jaguar | TEE Teeth | 7 | | | | | | | | Trade | A | Online marketplace |
| Chinese | Online marketplace | Online-marketplace-20 | OM-20 | 2019 | China | [NOT SPECIFI | [NOT DISPLA | [NOT DISPLA | TRANSFER SE | Jaguar | TEE Teeth | 7 | | | | | | | | Trade | A | Online marketplace |
| Chinese | Social network | Social-network-02 | SN-02 | 2019 | | [NOT SPECIFI | [NOT DISPLA | ADDITIONAL | [NOT DISPLA | Jaguar | TEE Teeth | [NOT SPECIFIED] | | | | | | | | Trade | [NO IMAGE REVIEW] | Social network |
| Chinese | Social network | Social-network-02 | SN-02 | 2018 | | [NOT SPECIFI | [NOT DISPLA | ADDITIONAL | [NOT DISPLA | Jaguar | TEE Teeth | 4 | | | | | | | | Trade | B | Social network |
| Chinese | Social network | Social-network-02 | SN-02 | 2018 | | [NOT SPECIFI | [NOT DISPLA | [NOT DISPLA | [NOT DISPLA | Jaguar | TEE Teeth | 1 | | | | | | | | Trade | B | Social network |
| Chinese | Social network | Social-network-02 | SN-02 | 2018 | | [NOT SPECIFI | [NOT DISPLA | ADDITIONAL | [NOT DISPLA | Jaguar | TEE Teeth | 11 | | | | | | | | Trade | B | Social network |
| Chinese | Social network | Social-network-02 | SN-02 | 2018 | | [NOT SPECIFI | [NOT DISPLA | ADDITIONAL | [NOT DISPLA | Jaguar | TEE Teeth | 3 | | | | | | | | Trade | B | Social network |
| Chinese | Social network | Social-network-02 | SN-02 | 2015 | | [NOT SPECIFI | DISPLAYED | ADDITIONAL | [NOT DISPLA | Jaguar | BPR Bone products | 1 | | | | | | | | Trade | B | Social network |
| Chinese | Social network | Social-network-02 | SN-02 | 2018 | | [NOT SPECIFI | [NOT DISPLA | [NOT DISPLA | [NOT DISPLA | Jaguar | TEE Teeth | 4 | | | | | | | | Trade | A | Social network |
| Chinese | Social network | Social-network-02 | SN-02 | 2018 | | [NOT SPECIFI | [NOT DISPLA | ADDITIONAL | [NOT DISPLA | Jaguar | TEE Teeth | 1 | | | | | | | | Trade | B | Social network |
| Chinese | Social network | Social-network-02 | SN-02 | 2017 | | [NOT SPECIFI | [NOT DISPLA | [NOT DISPLA | [NOT DISPLA | Jaguar | TEE Teeth | 6 | | | | | | | | Trade | A | Social network |
| Chinese | Social network | Social-network-02 | SN-02 | 2017 | | [NOT SPECIFI | [NOT DISPLA | ADDITIONAL | [NOT DISPLA | Jaguar | BOP Bone pieces | [NOT SPECIFIED] | | | | | | | | Trade | [NO IMAGE REVIEW] | Social network |
| Chinese | Social network | Social-network-02 | SN-02 | 2017 | | [NOT SPECIFI | [NOT DISPLA | ADDITIONAL | [NOT DISPLA | Jaguar | BOP Bone pieces | 1 | | | | | | | | Trade | B | Social network |
| Chinese | Social network | Social-network-02 | SN-02 | 2016 | | [NOT SPECIFI | [NOT DISPLA | [NOT DISPLA | [NOT DISPLA | Jaguar | SKI Skins | 2 | | Jaguar | WHI Whiske | [NOT SPECIFIED] | | | | Trade | A | Social network |
| Chinese | Social network | Social-network-02 | SN-02 | 2016 | | [NOT SPECIFI | [NOT DISPLA | ADDITIONAL | [NOT DISPLA | Jaguar | TEE Teeth | [NOT SPECIFIED] | | | | | | | | Trade | [NO IMAGE REVIEW] | Social network |
| Chinese | Social network | Social-network-02 | SN-02 | 2016 | | [NOT SPECIFI | [NOT DISPLA | ADDITIONAL | [NOT DISPLA | Jaguar | TEE Teeth | 4 | | | | | | | | Trade | A | Social network |
| Chinese | Social network | Social-network-02 | SN-02 | 2016 | | [NOT SPECIFI | [NOT DISPLA | [NOT DISPLA | [NOT DISPLA | Jaguar | TEE Teeth | 4 | | | | | | | | Trade | A | Social network |

| | | | | | | | | | | | | | | | | | | |
|---|---|---|---|---|---|---|---|---|---|---|---|---|---|---|---|---|---|---|
| Chinese | Social network | Social-network-02 | SN-02 | 2016 | | [NOT SPECIFI | [NOT DISPLA | ADDITIONAL | [NOT DISPLA | Jaguar | [NOT SPECIFIED] | [NOT SPECIFIED] | | | | Trade | [NO IMAGE REVIEW] | Social network |
| Chinese | Social network | Social-network-02 | SN-02 | 2016 | | [NOT SPECIFI | [NOT DISPLA | [NOT DISPLA | [NOT DISPLA | Jaguar | TEE Teeth | 8 | | | | Trade | [NO IMAGE REVIEW] | Social network |
| Chinese | Social network | Social-network-02 | SN-02 | 2016 | | [NOT SPECIFI | [NOT DISPLA | [NOT DISPLA | [NOT DISPLA | Jaguar | TEE Teeth | [NOT SPECIFIED] | | | | Trade | [NO IMAGE REVIEW] | Social network |
| Chinese | Social network | Social-network-02 | SN-02 | 2016 | | [NOT SPECIFI | [NOT DISPLA | [NOT DISPLA | [NOT DISPLA | Jaguar | TEE Teeth | 3 | | | | Trade | B | Social network |
| Chinese | Social network | Social-network-02 | SN-02 | 2016 | | [NOT SPECIFI | DISPLAYED | ADDITIONAL | [NOT DISPLA | Jaguar | TAI Tails | [NOT SPECIFIED BY RESEARCHER BUT IMAGE/S AVAILABLE] | | | | Trade | C | Social network |
| Chinese | Social network | Social-network-02 | SN-02 | 2016 | | [NOT SPECIFI | [NOT DISPLA | ADDITIONAL | [NOT DISPLA | Jaguar | CLA Claws | 1 | | | | Trade | B | Social network |
| Chinese | Social network | Social-network-02 | SN-02 | 2015 | | [NOT SPECIFI | [NOT DISPLA | ADDITIONAL | [NOT DISPLA | Jaguar | TEE Teeth | [NOT SPECIFIED] | | | | Trade | [NO IMAGE REVIEW] | Social network |
| Chinese | Social network | Social-network-02 | SN-02 | 2015 | | [NOT SPECIFI | [NOT DISPLA | [NOT DISPLA | [NOT DISPLA | Jaguar | TEE Teeth | 1 | | | | Trade | B | Social network |
| Chinese | Social network | Social-network-02 | SN-02 | 2015 | | [NOT SPECIFI | [NOT DISPLA | [NOT DISPLA | [NOT DISPLA | Jaguar | SKI Skins | 1 | | | | Trade | [NO IMAGE REVIEW] | Social network |
| Chinese | Social network | Social-network-02 | SN-02 | 2015 | China | [NOT SPECIFI | [NOT DISPLA | [NOT DISPLA | [NOT DISPLA | Jaguar | [NOT SPECIFIED] | [NOT SPECIFIED] | | | | Trade | [NO IMAGE REVIEW] | Social network |
| Chinese | Search engine | Online-marketplace-17 | OM-17 | 2015 | | [NOT SPECIFI | [NOT DISPLA | ADDITIONAL | TRANSFER SE | Jaguar | TEE Teeth | 4 | | | | Trade | C | Online marketplace |
| Chinese | Search engine | Online-marketplace-04 | OM-04 | 2011 | | [NOT SPECIFI | [NOT DISPLA | [NOT DISPLA | [NOT DISPLA | Jaguar | TEE Teeth | 1 | | | | Trade | A | Online marketplace |
| Chinese | Online marketplace | Online-marketplace-21 | OM-21 | [NOT SPECIFI | China | [NOT SPECIFI | DISPLAYED | [NOT DISPLA | [NOT DISPLA | Jaguar | TEE Teeth | 1 | | | | Trade | C | Online marketplace |
| Chinese | Social network | Social-network-03 | SN-03 | [NOT SPECIFIED] | | [NOT SPECIFI | [NOT DISPLA | [NOT DISPLA | [NOT DISPLA | Jaguar | TEE Teeth | 1 | | | | Trade | B | Social network |
| Chinese | Online marketplace | Online-marketplace-21 | OM-21 | [NOT SPECIFI | China | [NOT SPECIFI | [NOT DISPLA | [NOT DISPLA | [NOT DISPLA | Jaguar | TEE Teeth | 1 | | | | Trade | C | Online marketplace |
| Chinese | Online marketplace | Online-marketplace-21 | OM-21 | [NOT SPECIFI | China | [NOT SPECIFI | [NOT DISPLA | [NOT DISPLA | [NOT DISPLA | Jaguar | TEE Teeth | 1 | | | | Trade | C | Online marketplace |
| Chinese | Online marketplace | Online-marketplace-21 | OM-21 | [NOT SPECIFI | China | [NOT SPECIFI | [NOT DISPLA | [NOT DISPLA | [NOT DISPLA | Jaguar | TEE Teeth | 4 | | | | Trade | C | Online marketplace |
| Chinese | Online marketplace | Online-marketplace-18 | OM-18 | [NOT SPECIFI | China | [NOT SPECIFI | [NOT DISPLA | ADDITIONAL | TRANSFER SE | Jaguar | TEE Teeth | 2 | | | | Trade | A | Online marketplace |
| Chinese | Social network | Social-network-01 | SN-01 | 2017 | Peru/Ecuado | [NOT SPECIFI | [NOT DISPLA | [NOT DISPLA | [NOT DISPLA | Jaguar | TEE Teeth | 6 | | | | Trade | B | Social network |
| Chinese | Social network | Social-network-01 | SN-01 | 2016 | Peru/Ecuado | [NOT SPECIFI | [NOT DISPLA | [NOT DISPLA | [NOT DISPLA | Jaguar | TEE Teeth | 2 | | | | Trade | A | Social network |
| Chinese | Social network | Social-network-01 | SN-01 | 2016 | Peru/Ecuado | [NOT SPECIFI | [NOT DISPLA | [NOT DISPLA | [NOT DISPLA | Jaguar | BPR Bone products | [NOT SPECIFIED BY RESEARCHER BUT IMAGE/S AVAILABLE] | | | | Trade | B | Social network |
| Chinese | Social network | Social-network-01 | SN-01 | 2016 | Peru/Ecuado | [NOT SPECIFI | [NOT DISPLA | [NOT DISPLA | [NOT DISPLA | Jaguar | TEE Teeth | 2 | | | | Trade | B | Social network |
| Chinese | Social network | Social-network-01 | SN-01 | 2016 | Peru/Ecuado | [NOT SPECIFI | [NOT DISPLA | [NOT DISPLA | [NOT DISPLA | Jaguar | TEE Teeth | 1 | | | | Trade | A | Social network |
| Chinese | Social network | Social-network-01 | SN-01 | 2015 | Peru/Ecuado | [NOT SPECIFI | [NOT DISPLA | [NOT DISPLA | [NOT DISPLA | Jaguar | TEE Teeth | 2 | | | | Trade | B | Social network |
| Chinese | Social network | Social-network-01 | SN-01 | 2015 | Peru/Ecuado | [NOT SPECIFI | [NOT DISPLA | [NOT DISPLA | [NOT DISPLA | Jaguar | TEE Teeth | 1 | | | | Trade | A | Social network |
| Chinese | Social network | Social-network-01 | SN-01 | 2015 | Peru/Ecuado | [NOT SPECIFI | [NOT DISPLA | [NOT DISPLA | [NOT DISPLA | Jaguar | TEE Teeth | 1 | | | | Trade | B | Social network |
| Chinese | Social network | Social-network-01 | SN-01 | 2016 | Peru/Ecuado | [NOT SPECIFI | [NOT DISPLA | [NOT DISPLA | [NOT DISPLA | Jaguar | TEE Teeth | 1 | | | | Trade | B | Social network |
| Chinese | Social network | Social-network-01 | SN-01 | 2016 | Peru/Ecuado | [NOT SPECIFI | [NOT DISPLA | [NOT DISPLA | [NOT DISPLA | Jaguar | TEE Teeth | 3 | | | | Trade | C | Social network |
| Chinese | Social network | Social-network-01 | SN-01 | 2016 | Peru/Ecuado | [NOT SPECIFI | [NOT DISPLA | [NOT DISPLA | [NOT DISPLA | Jaguar | TEE Teeth | 2 | Jaguar | SKI Skins | [NOT SPECIFIED] | Trade | A | Social network |
| Chinese | Social network | Social-network-01 | SN-01 | 2016 | Peru/Ecuado | [NOT SPECIFI | [NOT DISPLA | [NOT DISPLA | [NOT DISPLA | Jaguar | TEE Teeth | 1 | | | | Trade | C | Social network |
| Chinese | Social network | Social-network-01 | SN-01 | 2016 | Peru/Ecuado | [NOT SPECIFI | [NOT DISPLA | [NOT DISPLA | [NOT DISPLA | Jaguar | TEE Teeth | 1 | | | | Trade | C | Social network |
| Chinese | Social network | Social-network-01 | SN-01 | 2016 | Peru/Ecuado | [NOT SPECIFI | [NOT DISPLA | [NOT DISPLA | [NOT DISPLA | Jaguar | TEE Teeth | 2 | | | | Trade | A | Social network |
| Chinese | Online marketplace | Online-marketplace-20 | OM-20 | 2019 | China | [NOT SPECIFI | [NOT DISPLA | [NOT DISPLA | TRANSFER SE | Jaguar | TEE Teeth | 2 | | | | Trade | A | Online marketplace |
| Chinese | Social network | Social-network-02 | SN-02 | 2019 | | [NOT SPECIFI | [NOT DISPLA | ADDITIONAL | [NOT DISPLA | Jaguar | TEE Teeth | 5 | | | | Trade | A | Social network |
| Chinese | Search engine | Online-marketplace-12 | OM-12 | [NOT SPECIFIED] | | [NOT SPECIFI | [NOT DISPLA | [NOT DISPLA | [NOT DISPLA | Jaguar | CLA Claws | [NOT SPECIFIED BY RESEARCHER BUT IMAGE/S AVAILABLE] | | | | Trade | C | Online marketplace |
| Chinese | Search engine | Online-marketplace-01 | OM-01 | [NOT SPECIFIED] | | [NOT SPECIFI | [NOT DISPLA | [NOT DISPLA | [NOT DISPLA | Jaguar | SKU Skulls | [NOT SPECIFIED] | | | | Trade | [NO IMAGE REVIEW] | Online marketplace |
| Chinese | Search engine | Search-engine-01 | SE-01 | 2010 | China | [NOT SPECIFI | [NOT DISPLA | [NOT DISPLA | [NOT DISPLA | Jaguar | LIV Live | 1 | | | | Trade | [NO IMAGE REVIEW] | Search engine |
| Chinese | Social network | Social-network-02 | SN-02 | 2019 | | [NOT SPECIFI | [NOT DISPLA | [NOT DISPLA | [NOT DISPLA | Jaguar | TEE Teeth | [NOT SPECIFIED] | Tiger | TEE Teeth | [NOT SPECIFIED] | Trade | [NO IMAGE REVIEW] | Social network |
| Chinese | Search engine | Social-network-02 | SN-02 | 2019 | | [NOT SPECIFI | [NOT DISPLA | [NOT DISPLA | [NOT DISPLA | Jaguar | TEE Teeth | [NOT SPECIFIED] | | | | Trade | [NO IMAGE REVIEW] | Social network |
| Chinese | Search engine | Social-network-02 | SN-02 | 2019 | | [NOT SPECIFI | [NOT DISPLA | [NOT DISPLA | [NOT DISPLA | Jaguar | TEE Teeth | 1 | | | | Trade | C | Social network |
| Chinese | Search engine | Social-network-02 | SN-02 | 2019 | | [NOT SPECIFI | [NOT DISPLA | [NOT DISPLA | [NOT DISPLA | Jaguar | TEE Teeth | [NOT SPECIFIED] | | | | Trade | [NO IMAGE REVIEW] | Social network |
| Chinese | Search engine | Social-network-02 | SN-02 | 2019 | | [NOT SPECIFI | [NOT DISPLA | [NOT DISPLA | [NOT DISPLA | Jaguar | TEE Teeth | [NOT SPECIFIED] | | | | Trade | [NO IMAGE REVIEW] | Social network |
| Chinese | Social network | Social-network-02 | SN-02 | 2019 | | [NOT SPECIFI | [NOT DISPLA | [NOT DISPLA | [NOT DISPLA | Jaguar | TEE Teeth | [NOT SPECIFIED] | | | | Trade | [NO IMAGE REVIEW] | Social network |
| Dutch | Online marketplace | Online-marketplace-19 | OM-19 | 2019 | Netherlands | BY POST | [NOT DISPLA | [NOT DISPLA | [NOT DISPLA | Jaguar | CAR Carvings (not ivo | 1 | | | | Trade | C | Online marketplace |
| Spanish | Social network | Social-network-01 | SN-01 | 2018 | Bolivia | [NOT SPECIFI | [NOT DISPLA | [NOT DISPLA | CONTACT SE | Jaguar | TEE Teeth | 2 | | | | Trade | [NO IMAGE REVIEW] | Social network |
| Spanish | Social network | Social-network-01 | SN-01 | 2018 | Bolivia | [NOT SPECIFI | DISPLAYED | [NOT DISPLA | [NOT DISPLA | Jaguar | HEA Heads | 1 | | | | Trade | A | Social network |
| Spanish | Social network | Social-network-01 | SN-01 | 2017 | Bolivia | [NOT SPECIFI | [NOT DISPLA | [NOT DISPLA | [NOT DISPLA | Jaguar | SKI Skins | 1 | | | | Trade | C | Social network |
| Spanish | Social network | Social-network-01 | SN-01 | 2018 | Bolivia | [NOT SPECIFI | [NOT DISPLA | [NOT DISPLA | CONTACT SE | Jaguar | SKI Skins | 1 | | | | Trade | [NO IMAGE REVIEW] | Social network |
| Spanish | Social network | Social-network-01 | SN-01 | 2018 | Bolivia | [NOT SPECIFI | DISPLAYED | [NOT DISPLA | CONTACT SE | Jaguar | SKI Skins | 1 | | | | Trade | A | Social network |
| Spanish | Social network | Social-network-01 | SN-01 | 2018 | Bolivia | [NOT SPECIFI | [NOT DISPLA | [NOT DISPLA | [NOT DISPLA | Jaguar | TEE Teeth | 1 | | | | Trade | A | Social network |
| Spanish | Social network | Social-network-01 | SN-01 | 2018 | Bolivia | [NOT SPECIFI | DISPLAYED | [NOT DISPLA | CONTACT SE | Jaguar | HEA Heads | 1 | | | | Trade | A | Social network |
| Spanish | Social network | Social-network-01 | SN-01 | 2018 | Bolivia | [NOT SPECIFI | [NOT DISPLA | [NOT DISPLA | [NOT DISPLA | Jaguar | SKI Skins | 1 | | | | Trade | [NO IMAGE REVIEW] | Social network |
| Spanish | Social network | Social-network-01 | SN-01 | 2017 | Bolivia | [NOT SPECIFI | DISPLAYED | [NOT DISPLA | CONTACT SE | Jaguar | TEE Teeth | 8 | | | | Trade | A | Social network |
| Spanish | Social network | Social-network-01 | SN-01 | 2018 | Bolivia | [NOT SPECIFI | DISPLAYED | [NOT DISPLA | CONTACT SE | Jaguar | HEA Heads | 1 | | | | Trade | A | Social network |
| Spanish | Social network | Social-network-01 | SN-01 | 2018 | Bolivia | [NOT SPECIFI | DISPLAYED | [NOT DISPLA | CONTACT SE | Jaguar | HEA Heads | 1 | | | | Trade | A | Social network |
| Spanish | Social network | Social-network-01 | SN-01 | 2018 | Bolivia | [NOT SPECIFI | DISPLAYED | [NOT DISPLA | CONTACT SE | Jaguar | TEE Teeth | 2 | | | | Trade | A | Social network |
| Spanish | Social network | Social-network-01 | SN-01 | 2019 | Bolivia | [NOT SPECIFI | DISPLAYED | [NOT DISPLA | CONTACT SE | Jaguar | SKI Skins | 1 | | | | Trade | [NO IMAGE REVIEW] | Social network |
| Spanish | Social network | Social-network-01 | SN-01 | 2019 | Bolivia | [NOT SPECIFI | [NOT DISPLA | [NOT DISPLA | CONTACT SE | Jaguar | HEA Heads | 1 | Jaguar | SKI Skins | 1 | Trade | [NO IMAGE REVIEW] | Social network |
| Spanish | Social network | Social-network-01 | SN-01 | 2019 | Bolivia | [NOT SPECIFI | DISPLAYED | ADDRESS DIS | [NOT DISPLA | Jaguar | OIL Oil | 0.25 ml | | | | Trade | B | Social network |
| Spanish | Social network | Social-network-01 | SN-01 | 2019 | Bolivia | [NOT SPECIFI | [NOT DISPLA | [NOT DISPLA | [NOT DISPLA | Jaguar | SKI Skins | 1 | | | | Trade | A | Social network |
| Spanish | Social network | Social-network-01 | SN-01 | 2018 | Bolivia | [NOT SPECIFI | DISPLAYED | [NOT DISPLA | [NOT DISPLA | Jaguar | TEE Teeth | 1 | | | | Trade | [NO IMAGE REVIEW] | Social network |
| Spanish | Social network | Social-network-01 | SN-01 | 2019 | Bolivia | [NOT SPECIFI | DISPLAYED | [NOT DISPLA | [NOT DISPLA | Jaguar | HEA Heads | 1 | Jaguar | SKI Skins | 1 | Trade | [NO IMAGE REVIEW] | Social network |
| Spanish | Social network | Social-network-01 | SN-01 | 2018 | Bolivia | [NOT SPECIFI | DISPLAYED | [NOT DISPLA | [NOT DISPLA | Jaguar | HEA Heads | 1 | | | | Trade | [NO IMAGE REVIEW] | Social network |
| Spanish | Social network | Social-network-01 | SN-01 | 2018 | Bolivia | [NOT SPECIFI | DISPLAYED | [NOT DISPLA | [NOT DISPLA | Jaguar | TEE Teeth | 4 | | | | Trade | [NO IMAGE REVIEW] | Social network |
| Spanish | Social network | Social-network-01 | SN-01 | 2018 | Bolivia | [NOT SPECIFI | [NOT DISPLA | [NOT DISPLA | [NOT DISPLA | Jaguar | TEE Teeth | 1 | | | | Trade | A | Social network |
| Spanish | Social network | Social-network-01 | SN-01 | 2017 | Bolivia | [NOT SPECIFI | DISPLAYED | [NOT DISPLA | CONTACT SE | Jaguar | SKI Skins | 1 | | | | Trade | A | Social network |
| Spanish | Social network | Social-network-01 | SN-01 | 2017 | Bolivia | [NOT SPECIFI | [NOT DISPLA | [NOT DISPLA | [NOT DISPLA | Jaguar | TEE Teeth | 1 | | | | Trade | [NO IMAGE REVIEW] | Social network |
| Spanish | Social network | Social-network-01 | SN-01 | 2018 | Bolivia | [NOT SPECIFI | [NOT DISPLA | [NOT DISPLA | CONTACT SE | Jaguar | HEA Heads | 1 | | | | Trade | [NO IMAGE REVIEW] | Social network |
| Spanish | Social network | Social-network-01 | SN-01 | 2018 | Colombia | [NOT SPECIFI | DISPLAYED | [NOT DISPLA | [NOT DISPLA | Jaguar | TEE Teeth | 1 | | | | Trade | B | Social network |
| Spanish | Social network | Social-network-01 | SN-01 | 2009 | Colombia | [NOT SPECIFI | DISPLAYED | [NOT DISPLA | [NOT DISPLA | Jaguar | TEE Teeth | 1 | | | | Trade | [NO IMAGE REVIEW] | Social network |
| Spanish | Online marketplace | Online-marketplace-16 | OM-16 | 2017 | Ecuador | [NOT SPECIFI | [NOT DISPLA | EMAIL CONT | [NOT DISPLA | Jaguar | LIV Live | [NOT SPECIFIED] | | | | Trade | [NO IMAGE REVIEW] | Online marketplace |
| English | Social network | Social-network-01 | SN-01 | 2011 | Mexico | [NOT SPECIFI | [NOT DISPLA | [NOT DISPLA | [NOT DISPLA | Jaguar | TEE Teeth | 2 | | | | Trade | B | Social network |
