## Supplementary material for "Multi-lingual multi-platform investigations of online trade in jaguar parts": S3 - Effort log template

| Reference number for search (per individual searcher) | Date column, consistent formatting throughout | Person initials | Countries | Region | Language | Time search | Search terms used | Online platform/search engine | Categorisation of online platform/search engine (code for analysis) | Type of post | # productive returns | # unrelated returns | Total returns | Analysis Percentage of returns which were productive in finding posts for consideration in raw results, by search session and search string | Analysis Efficiency rate: productive returns by time spent per search session |
|---|---|---|---|---|---|---|---|---|---|---|---|---|---|---|---|
