## Supplementary material for "Multi-lingual multi-platform investigations of online trade in jaguar parts": S4 - Guidance - Standardization and identification of efficient search term & platform combinations

**S4. Guidance standardization & identification of most efficient search term & platform combinations**

When an investigation involves a variable number of searchers among the languages and within that, variable levels of effort expended by each individual searcher, crude summaries that rank frequency of jaguar posts by language will be skewed/biased by uneven effort expended across languages. Crude rankings of jaguar post prevalence across languages can be adjusted by the relative effort across the languages. In our investigation, the crude rank of prevalence of jaguar posts in each language was adjusted by: 1) time spent searching per language; and 2) number of searches in each language – to produce a less biased ranking of prevalence of jaguar posts in the respective languages.

Standardization was facilitated by a record of individual search effort in terms of time spent searching and in terms of numbers of searches with each search term/platform combination (***see Supporting File S3 Effort Log***). The total number of returns (posts) was examined, and within that, the number of posts with jaguar trade. Records of effort made by each researcher in each language, were used to obtain less biased estimates of relative frequency of jaguar trade posts per language, and to examine which search term/platform combinations were the most efficient.

Each researcher recorded effort by time, and by searches, ***linked to search term(s) and platform combinations*****,** and in the same row, total returns (posts with or without jaguar trade) and productive returns (posts with jaguar trade). ***See supporting file S3*** **Effort Log**.

We adjusted the gross ranking of productive returns (jaguar posts): 1) by effort (time, searches) for each/all language; yielding 2) a ranking of frequency of jaguar posts by each language adjusted by the relative effort (by time, by number of searches). These rankings provided a less biased perspective on the respective prevalence of jaguar posts across the languages that we searched.

By linking each investigator's search effort and productivity to search term(s)/platform combinations it was possible extract the most effective search term/platform combinations to detect illegal jaguar/wildlife trade. Using S3, we were able to evaluate productivity of each search term (s)/platform combinations. **We have decided to not publish that raw data for reasons that follow.**

While, relative productivity of search term/platform combinations: 1) could guide future research; 2) increase efficiency in detecting online trade in jaguar parts, and other illegal wildlife trade; 3) **publishing those analyses could facilitate evasion by traders, so we are not presenting those raw data files.**

We do present the relative ranking of posts of jaguar trade across the languages, obtained by adjusting the crude results by relative effort expended in each language (it is in the text and see Supporting File S3) for an effort-adjusted ranking of prevalence of jaguar trade posts across the respective languages.

We encourage future investigations to record effort, and productivity by time, searches, and search term(s)/platform combinations so that results can be interpreted with reduced sampling bias, and to increase efficiencies in detecting illegal trade in jaguar parts and other wildlife.
