## Supplementary material for "Multi-lingual multi-platform investigations of online trade in jaguar parts": S5 - Internet penetration

S5. Internet penetration, number of internet users and percentage of internet users per country (source: Miniwatts Marketing Group 2020).

| Country | Internet penetration (% total population using internet) | Number of internet users | Number of Facebook users | % users per country considering the total of users in Central and South America |
|---|---|---|---|---|
| Argentina | 0.931 | 41,586,960 | 30,000,000 | 9.9250 |
| Paraguay | 0.896 | 6,177,748 | 3,300,000 | 1.4743 |
| Uruguay | 0.882 | 3,059,727 | 2,400,000 | 0.7302 |
| Costa Rica | 0.859 | 4,296,443 | 3,200,000 | 1.0253 |
| Ecuador | 0.799 | 13,476,687 | 10,000,000 | 3.2163 |
| Chile | 0.775 | 14,108,392 | 13,000,000 | 3.3670 |
| Brazil | 0.707 | 149,057,635 | 139,000,000 | 35.5735 |
| Panama | 0.686 | 2,899,892 | 2,000,000 | 0.6920 |
| Peru | 0.676 | 22,000,000 | 20,000,000 | 5.2504 |
| Bolivia | 0.675 | 7,570,580 | 6,100,000 | 1.8067 |
| Mexico | 0.665 | 88,000,000 | 79,000,000 | 21.0017 |
| Colombia | 0.632 | 31,275,567 | 29,000,000 | 7.4641 |
| Suriname | 0.598 | 340,000 | 310,000 | 0.0811 |
| El Salvador | 0.574 | 3,700,000 | 3,400,000 | 0.8830 |
| Venezuela | 0.531 | 17,178,743 | 13,000,000 | 4.0998 |
| Belize | 0.513 | 200,020 | 200,000 | 0.0477 |
| Guyana | 0.505 | 395,007 | 360,000 | 0.0942 |
| Nicaragua | 0.425 | 2,700,000 | 2,500,000 | 0.6443 |
| Guatemala | 0.414 | 7,268,597 | 6,800,000 | 1.7346 |
| French Guiana | 0.414 | 120,000 | 110,000 | 0.0286 |
| Honduras | 0.376 | 3,600,000 | 3,400,000 | 0.8591 |
